# The ParB clamp docks onto Smc for DNA loading via a joint-ParB interface

**DOI:** 10.1101/2021.12.15.472096

**Authors:** Florian P. Bock, Anna Anchimiuk, Marie-Laure Diebold-Durand, Stephan Gruber

**Affiliations:** Department of Fundamental Microbiology (DMF), Faculty of Biology and Medicine (FBM), University of Lausanne, 1015 Lausanne, Switzerland

**Keywords:** Smc, condensin, DNA loop extrusion, ParB, ParABS, chromosome segregation

## Abstract

Chromosomes readily unlink from one another and segregate to daughter cells during cell division highlighting a remarkable ability of cells to organize long DNA molecules. SMC complexes mediate chromosome folding by DNA loop extrusion. In most bacteria, SMC complexes start loop extrusion at the ParB/*parS* partition complex formed near the replication origin. Whether they are recruited by recognizing a specific DNA structure in the partition complex or a protein component is unknown. By replacing genes in *Bacillus subtilis* with orthologous sequences from *Streptococcus pneumoniae*, we show that the three subunits of the bacterial Smc complex together with the ParB protein form a functional module that can organize and segregate chromosomes when transplanted into another organism. Using chimeric proteins and chemical cross-linking, we find that ParB binds to the Smc subunit directly. We map a binding interface to the Smc joint and the ParB CTP-binding domain. Structure prediction indicates how the ParB clamp presents DNA to the Smc complex to initiate DNA loop extrusion.

## Introduction

Organizing DNA for chromosome segregation is a fundamental challenge across all domains of life. Structural maintenance of chromosomes (SMC) complexes fold DNA into a loop or an arrangement of multiple loops via a process dubbed DNA loop extrusion (Yatskevich et al., 2019). They are ubiquitously found in the domains of life. Three types of SMC complexes with dedicated functions (*i.e*. Smc5/6, cohesin and condensin) are nearly universal in eukaryotes (Yoshinaga and Inagaki, 2021). In prokaryotes, the Smc-ScpAB complex is predominant—being widely distributed in bacteria and present in some archaea. Disruption of any of the three subunits of Smc-ScpAB results in a strong chromosome segregation defect in *B. subtilis*, ultimately leading to cell death when cells are grown on nutrient-rich medium that promotes fast DNA replication (Gruber et al., 2014). The other bacterial SMC variants, MukBEF and MksBEF(G), are highly diverged with the latter supporting plasmid restriction rather than chromosome segregation in some bacteria (Panas et al., 2014; Petrushenko et al., 2011).

The Smc-ScpAB complex is recruited to the replication origin region of the bacterial chromosome by 16 bp palindromic ‘*parS*’ DNA sequences that associate with the ParB protein (Gruber and Errington, 2009; Sullivan et al., 2009). Smc-ScpAB then starts translocating onto *parS*-flanking DNA in both orientations (Figure 1A). This bidirectional DNA translocation (*i.e*. DNA-loop-extrusion) brings together loci distantly located on opposing chromosome arms (Tran et al., 2017; Wang et al., 2017), which is thought to help separate the nascent sister chromosomes, presumably by removing DNA entanglements in the wake of the DNA replication forks (Bürmann and Gruber, 2015). How Smc-ScpAB recognizes the *parS* loading site, how it loads onto DNA, and how it starts loop extrusion is poorly understood.

**Figure 1.**
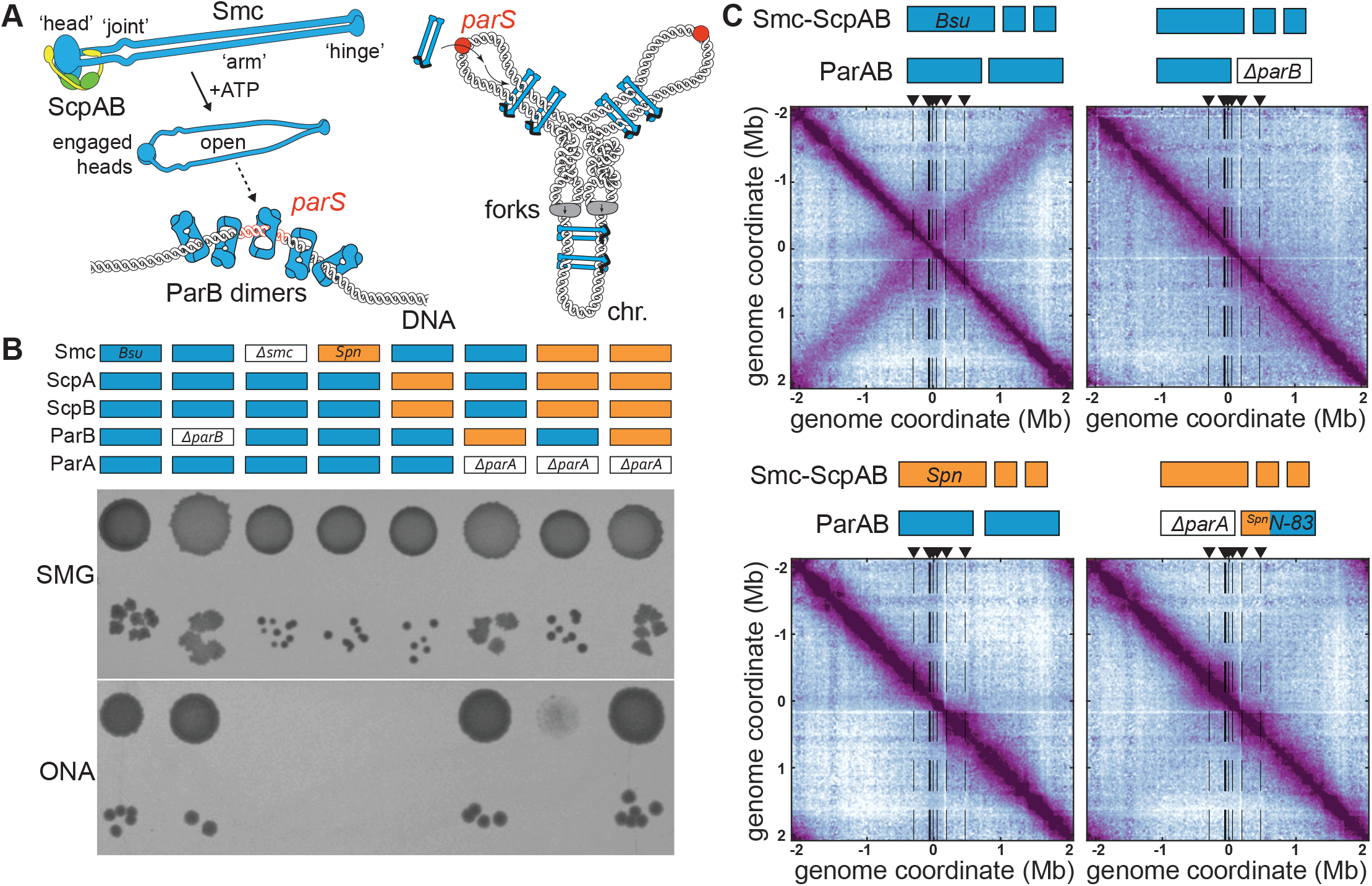
A four-component system for chromosome organization in bacteria. **A)** Schematic of Smc recruitment via ParB at *parS* sites (left panels) and chromosome organization of DNA loop extrusion (right panel). Chromosome, chr.; DNA replication forks, forks. **B)** Viability assessment of gene-transplanted strains by spotting on nutrient-poor medium (SMG) and nutrient-rich medium (ONA). Gene identity of strains indicated by colored bars, *Bsu* in blue colors, *Spn* in orange colors. **C)** Normalized 3C-seq contact maps of strains with indicated genotypes from exponentially growing cultures. Additional maps are shown in Supplementary Figure 1. All 3C-seq contact maps presented are divided into 10 kb bins. The replication origin is placed in the middle. The interaction score is in log10 scale (for more details go to Materials and Methods). Note that the contact map for wild type is same as in (Anchimiuk *et al*., 2021).

ParABS systems promote the partitioning of bacterial chromosomes and the stable maintenance of low copy-number plasmids. They comprise the *parS* sites, the ParA ATPase, and the ParB CTPase. ParB protein locally enriches in a ‘partition complex’ by binding the cofactor CTP and clamping onto *parS* DNA (**Figure 1A**). The clamps then spread onto flanking DNA by 1D diffusion and recruit further ParB dimers, before eventually unloading from the chromosome upon CTP hydrolysis (Antar et al., 2021; Jalal et al., 2021; Osorio-Valeriano et al., 2021; Soh et al., 2019; Tišma et al., 2021). ParB comprises three globular domains (**Figure 2B**). The amino-terminal “N domain” harbors the CTP binding pocket. It homodimerizes upon contact with *parS* DNA thus closing the ParB clamp. The middle “M domain” includes a helix-turn-helix motif which specifically recognizes *parS* DNA (Chen et al., 2015). The carboxy-terminal “C domain” serves to dimerize two ParB monomers and also promotes sequences-nonspecific DNA binding (Fisher et al., 2017; Schumacher and Funnell, 2005). ParA forms gradients on the bacterial chromosome along which the partition complexes move to equiposition themselves (Hwang et al., 2013; Lim et al., 2014). This requires stimulation of ParA ATP hydrolysis by an N-terminal peptide on ParB, which converts DNA-bound ParA dimers into cytosolic monomers (Gruber and Errington, 2009; Scholefield et al., 2011; Zhang and Schumacher, 2017).

**Figure 2.**
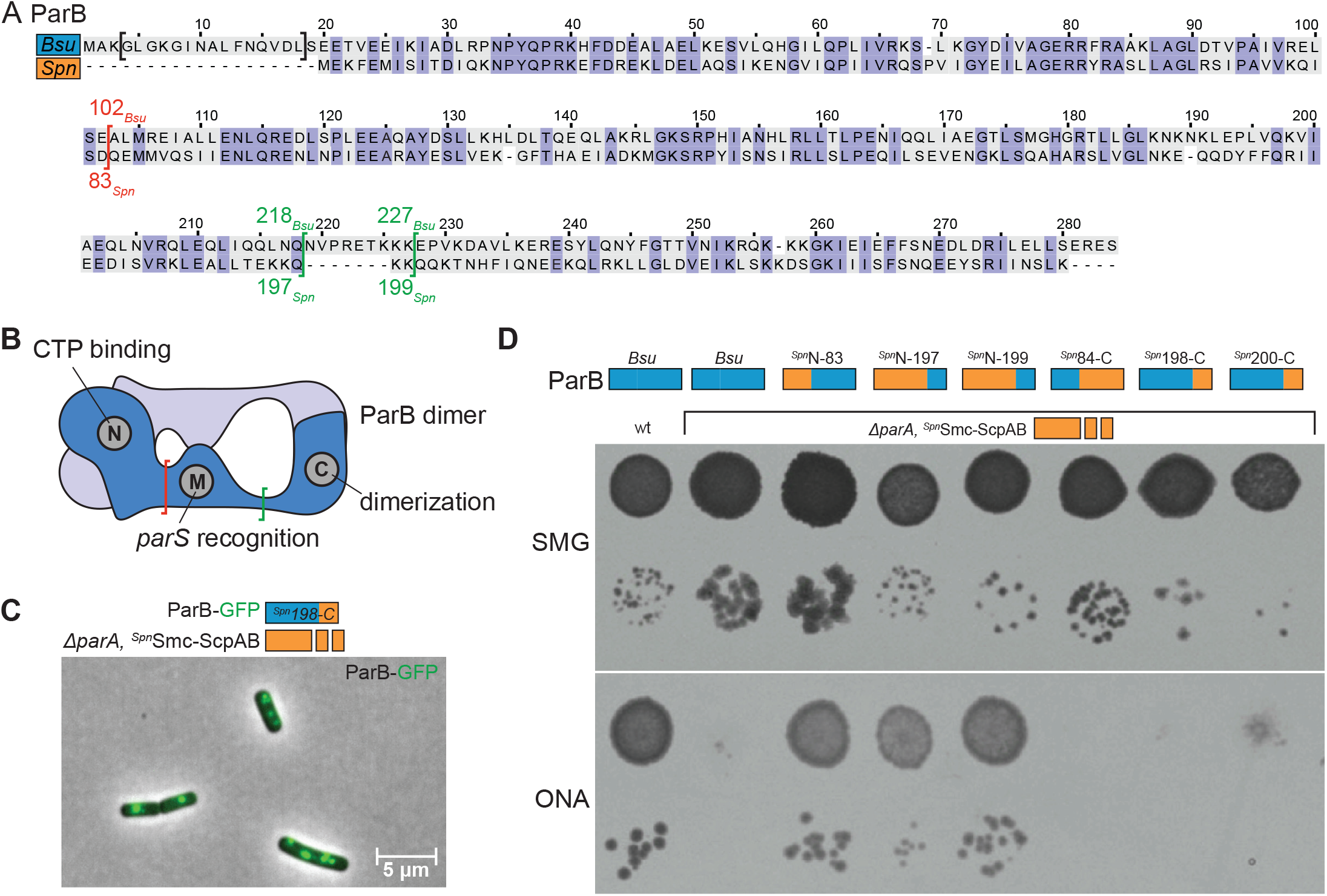
Smc binds the ParB N domain. **A)** Sequence alignment of *Bsu* and *Spn* ParB protein sequences. Identical residues denoted by blue background colors, divergent residues in grey colors. Construction of ParB chimeras is indicated in red colors at the N to M domain transition and in green colors for the M to C domain transition. **B)** ParB domain structure. Constructions of chimeras are indicated by brackets. **C)** Microscopy image of *B. subtilis* cells harbouring the ^*Spn*^Smc-ScpAB with the ^*Spn*^198-C ParB chimera fused to GFP protein. **D)** Spotting assay of *B. subtilis* strains carrying the ^*Spn*^Smc-ScpAB as well as the indicated chimeric ParB proteins. As in Figure 1B.

The Smc protein folds into a highly elongated particle having an ABC-type ATPase “head” domain at one end and a dimerization “hinge” domain at the other end of a long intramolecular antiparallel coiled-coil “arm” (**Figure 1A**) (Haering et al., 2002). The kleisin protein ScpA connects the head domain of one Smc subunit to the head-proximal arm (“neck”) of the other, together forming a ring-shaped protein complex capable of entrapping chromosomal DNA (Bürmann et al., 2013; Gligoris et al., 2014; Wilhelm et al., 2015). Two ScpB proteins—belonging to the kite family—bind to the central region of ScpA (Palecek and Gruber, 2015). The two long arms co-align to form a rod-shaped particle with mis-aligned head domains. ATP-engagement of the head domains in turn pulls the arms apart, thus creating a more open ring-shaped particle (Vazquez Nunez et al., 2021) (**Figure 1A**). An essential DNA binding interface is formed by ATP-engaged Smc head domains. How DNA is clamped at the Smc heads, and how DNA binding and ATP hydrolysis promote loop extrusion is not well understood (Vazquez Nunez et al., 2019).

Recruitment of Smc-ScpAB by the partition complex relies on several factors. The Smc heads have to bind ATP and engage with one another, while ATP hydrolysis by Smc is dispensable (Minnen et al., 2016). The hydrolysis-defective mutant Smc(E1118Q) (“EQ”) efficiently targets to *parS* DNA, especially when arm alignment is artificially weakened (e.g. by mutations preventing hinge dimerization). The accumulation of Smc(EQ) at *parS* DNA requires DNA clamping by ATP-engaged Smc heads (Vazquez Nunez *et al*., 2019). DNA clamping by ParB is also essential for Smc recruitment, while CTP hydrolysis is dispensable (Antar *et al*., 2021). Altogether, this suggests that an open, ATP-bound, DNA-clamping state of Smc-ScpAB associates with a ParB DNA sliding clamp. The interface between Smc-ScpAB and ParB however has remained elusive, possibly owing to a weak and transient nature of the interaction or the dependence on a cofactor. Also, the relationship of DNA in the ParB clamp and the Smc clamp is unclear.

Here we provide conclusive evidence that ParABS promotes chromosome folding via a direct protein-protein interaction between the ParB protein and the Smc-ScpAB complex. Using chimeric proteins and site-specific *in vivo* crosslinking we identify the key residues for specifying the interaction. These residues are located on the Smc joint and the recently discovered CTP-controlled DNA-gate domain of ParB (Antar *et al*., 2021; Soh *et al*., 2019). Structure prediction provides detailed insights into how the ParB clamp feeds DNA into the Smc-ScpAB complex for loop extrusion. We furthermore demonstrate that the Smc-ScpAB and ParB/*parS* complexes together form a minimal system for chromosome folding and segregation that can be transplanted from one bacterial species to another.

## Results

### A four-gene module from *S. pneumoniae* promotes chromosome segregation in *B. subtilis*

To determine the minimal set of factors needed to organize and segregate chromosomes in bacteria, we replaced genes encoding for components of the *B. subtilis* (*Bsu*) Smc holo-complex for orthologous counterparts. As gene donor, we chose *S. pneumoniae* (*Spn*) which belongs to the same branch of Gram-positive bacteria (the firmicutes) and also relies on Smc-ScpAB for chromosome segregation (Minnen et al., 2011). The Smc genes for example display only 38 % amino acid sequence identity, showing that the two species have substantially diverged and implying that the respective interaction partners significantly co-evolved. Substituting the *scpAB* operon, which encodes for the ScpA and ScpB subunits, or the *smc* gene by the respective *Spn* orthologs lead to severe growth defects on nutrient rich medium similar to the *Δsmc* mutant (**Figure 1B**). Combining the *smc* and *scpAB* genes of *Spn* origin only marginally improved growth on nutrient-rich medium, demonstrating that the genes encoding for the subunits of the *Spn* Smc complex alone or in combination (‘^*Spn*^Smc-ScpAB’) are unable to support chromosome segregation in *B. subtilis*. Chromosome folding was also altered in the ^Spn^Smc-ScpAB strain as judged from 3C-Seq contact maps (**Figure 1C**). Like in the *ΔparB* mutant, contacts along the secondary diagonal originating from arm-arm co-alignment by Smc-ScpAB are missing (**Figure 1C**)(Wang *et al*., 2017). ^*Bsu*^Smc-ScpAB however loads onto the chromosome sporadically even in the absence of ParB, leading to residual contacts across the chromosome arms, which are widely distributed in the inter-arm (top-right and bottom-left) quadrants of the *ΔparB* contact map. Residual loop extrusion activity by ^*Bsu*^Smc-ScpAB starting from random positions may thus support chromosome segregation and cell viability in the *ΔparB* strain. ^*Spn*^Smc-ScpAB apparently fails to productively load onto the chromosome at *parS* or elsewhere (as evident from the clear reduction of inter-arm contacts) and is thus unable to support chromosome segregation.

One explanation for the strong phenotypes associated with ^*Spn*^Smc-ScpAB might be its inability to interact with one or more host factors in *B. subtilis*. Known factors include the ParB protein and the *parS* sites—that together form the partition complex targeting Smc-ScpAB to the replication origin region in *B. subtilis* and in *S. pneumoniae* (Gruber and Errington, 2009; Minnen *et al*., 2011; Sullivan *et al*., 2009). While the function of endogenous ^*Bsu*^Smc-ScpAB does not strictly require ParB or *parS, ^Spn^*Smc-ScpAB may rely on cognate ParB even for basal functions in *B. subtilis*. To test this possibility, we next substituted the *parB* gene. We added a *Spn parS* site at the 3’ end of *Spn parB* (**Supplementary Figure 2A**) because the *Bsu parB* gene harbours an internal *parS* sequence (Minnen *et al*., 2011). *S. pneumoniae* does not encode for ParA, and its ParB sequence lacks the N-terminal extension that normally stimulates ATP hydrolysis by ParA (Gruber and Errington, 2009; Leonard et al., 2005). To eliminate any detrimental effect by unregulated *Bsu* ParA (Murray and Errington, 2008; Quisel and Grossman, 2000), we thus excluded the first 20 amino acids of the *Bsu parB* gene from any *Bsu/Spn* chimeric ParB sequences and also deleted the neighboring *parA* gene. As expected from the weak growth phenotypes of Δ*parB*, these modifications on their own did not noticeably alter cell viability (but caused a change in colony morphology) (**Figure 1B**). Introducing *Spn* ParB into strains already harboring *Spn* Smc and *Spn* ScpAB resulted in much improved growth on nutrient-rich medium with the viability and growth being comparable to wild-type cells (**Figure 1B**). Likewise, 3C-Seq analysis showed increased levels of contacts across the left and the right chromosome arm (when relevant parts of ParB comprised the *Spn* sequence; see below) (**Figure 1C**). The levels of these contacts were still reduced when compared to wild type, and their distribution was broadened implying that ^Spn^Smc-ScpAB is less efficient or less organized in forming these contacts (with all *parS* sites or only two *parS* sites present) (Anchimiuk et al., 2021). Nevertheless, these results demonstrate that ^*Spn*^Smc-ScpAB is principally capable of organizing the chromosome for efficient segregation in *B. subtilis*, but only when being targeted to the replication origin region by its cognate ^*Spn*^ParB.

These findings show that Smc-ScpAB collaborates only with ParABS to organize chromosomes in two distantly related bacteria. Functional interactions of four proteins—ParB, Smc, ScpA and ScpB—are needed for proper chromosome folding. Sequence divergence prevents productive protein-protein interaction across the two species rendering the *Bsu* and *Spn* modules orthogonal to one another. The critical involvement of other host proteins in chromosome folding by Smc is moreover highly unlikely, as such factors would have to fruitfully interact with *Bsu* and *Spn* proteins despite their divergent sequences.

### Smc binds to the N-terminal CTP-binding domain of ParB

We next made use of the orthogonality of the chromosome folding modules to map protein binding interfaces between ParB and Smc-ScpAB. We engineered chimeric proteins having either amino- or carboxy-terminal *Bsu* ParB sequences exchanged for the corresponding *Spn* sequence (**Figure 2A**). The junctions of the chimeric proteins were chosen in regions of low sequence conservation and at domain boundaries to try to minimize protein folding problems (**Figure 2B**). A GFP-tagged variant of the ParB chimera ^*Spn*^198-C showed clear focal localization suggesting that it self-loads onto *parS* DNA efficiently (**Figure 2C**, **Supplementary Figure 2B**) (Glaser et al., 1997). The GFP-tagged versions of chimera ^*Spn*^N-197 and of full-length *Spn* ParB also displayed focal localization, albeit with the foci being more diffuse as previously observed with ParB-GFP in *S. pneumoniae* (**Supplementary Figure 2C**) (Kjos and Veening, 2014; Minnen *et al*., 2011). As expected, none of the chimeric ParB proteins led to obvious growth phenotypes in otherwise wild-type strains. When introduced into a ^*Spn*^Smc-ScpAB strain, however, all chimeric ParB proteins with amino-terminal regions of *Spn* origin (^*Spn*^N-83, ^*Spn*^N-197 and ^*Spn*^N-199) promoted robust growth on nutrient-rich medium (**Figure 2D**). When instead the ParB carboxy-terminus originated from *S. pneumoniae* (^*Spn*^84-C, ^*Spn*^198-C and ^*Spn*^200-C) the strains showed poor growth on nutrient-rich medium as with full-length *Bsu* ParB (**Figure 2D**). These observations strongly suggest that the CTP-binding domain of ParB is responsible for a direct physical and functional interaction with Smc-ScpAB. The 3C-Seq analysis mentioned above is consistent with this notion as it showed increased trans-chromosome arm contacts in ^Spn^Smc-ScpAB when ^*Spn*^N-83 was also present (**Figure 1C**).

### Identifying ParB residues critical for Smc association

To elucidate how Smc and ParB may orient one another, we next set out to fine-map the Smc binding interface on ParB by identifying which *Spn* ParB residues are required to support the action of ^Spn^Smc-ScpAB in *B. subtilis*. There are thirty-eight residue differences across the relevant amino-terminal part of *Bsu* and *Spn* ParB. We focused on surface-exposed residues and grouped them into four patches for mutagenesis (denoted as 1 to 4) (**Figure 3A**). Exchange of all four patches in *Bsu* ParB led to robust growth of the ^*Spn*^Smc-ScpAB strain (**Figure 3B**) comparable to full exchange of the amino-terminal sequence (^*Spn*^N-83), indicating that the residue differences (8 in total) outside the chosen patches are not critically relevant. Strains harboring *Spn* residues only in three of the four patches should exhibit different growth behaviors: A strain harboring *Bsu* residues in patch 1 or in patch 3 failed to grow on nutrient-rich medium, highlighting their importance for ParB-Smc interaction (**Figure 3B**). A strain harboring *Bsu* residues only in patch 2 showed good growth, while a strain with a *Bsu* patch 4 displayed an intermediate phenotype indicating that patch 4 is somewhat important while patch 2 is largely dispensable. We conclude that residues from patches 1, 3, and 4 are noticeably contributing to the interaction with Smc.

**Figure 3.**
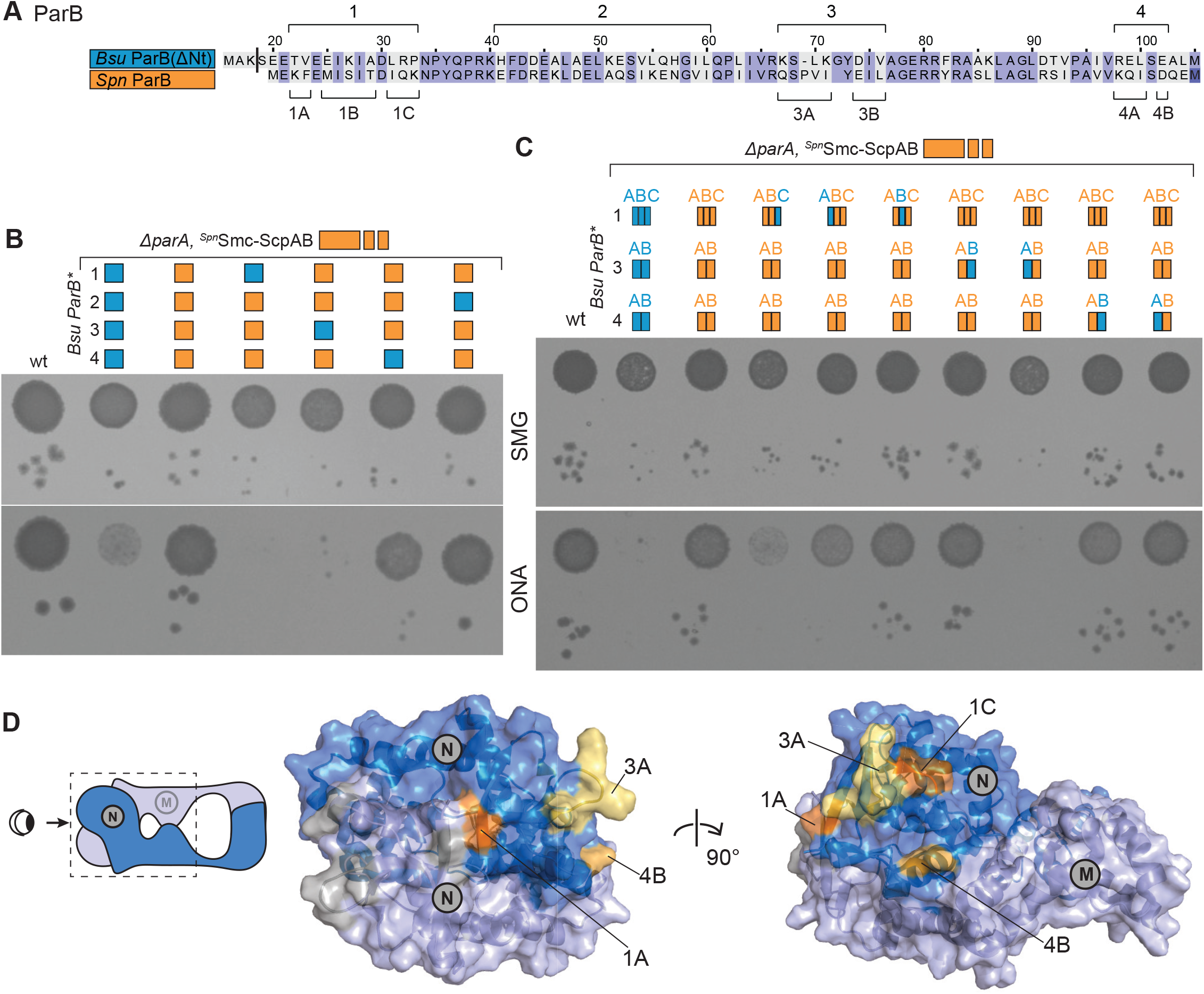
Fine mapping of the Smc binding site on ParB. **A)** Grouping of ParB residues in patches (1 to 4, top labels) and sub-patches (A, B and C, bottom labels). Sequence alignment of *Bsu* and *Spn* ParB N domain. As in Figure 2A. **B)** Viability assay by dilution spotting for ^*Spn*^Smc-ScpAB strains carrying chimeric ParB proteins as given in A). As in Figure 1B. **C)** As in Figure 3B for ParB chimeras with sub-patches. **D)** Distribution of the identified Smc-interacting residues on the CTP engaged ParB N-domain dimer in surface plot representation (PDB: 6SDK, (Soh *et al*., 2019)). ParB chains are shown in blue and grey colors, respectively. Key residues are indicated and highlighted in yellow, orange, and brown colors. Notably, the presence of *parA* significantly reduced the viability of some of the strains, indicating that ParA mis-regulation is indeed not tolerated well by these ParB variants in combination with ^*Spn*^Smc-ScpAB (**Supplementary Figure 3A**).

Following the same strategy, we sub-divided these residues into patches 1A, 1B, 1C, 3A, 3B, 4A, and 4B (**Figure 3A**). Converting these patches individually to the *Bsu* sequence demonstrated that residues in 1B, 3B, 4A, and 4B are largely dispensable for promoting Smc-ParB interaction. On the contrary, residues in patches 1A, 1C, and 3A appear critical (**Figure 3C**) with the conversion of patch 3A having a particularly severe impact on growth. 1A, 1C, and 3A together comprise eight residue differences, which mapped closely together on the surface of the ParB-CDP crystal structure (PDB: 6SDK) (Soh *et al*., 2019), together delineating a putative Smc-binding interface on ParB (**Figure 3D**). The immediate proximity of the identified putative binding interface to the CTP-binding pocket supports the notion that chromosomal loading of Smc is closely linked to ParB CTP binding and hydrolysis and thus potentially coupled to other cellular activities including DNA replication initiation and ParABS segregation (Antar *et al*., 2021). Of note, the chimeric ParB proteins (with one exception) performed well in an Smc-pk3 strain (Gruber and Errington, 2009), that is sensitized for ParB function by the hypomorphic *smc* allele. This indicates that the chimeric proteins are able to support *Bsu* Smc-ScpAB function, presumably by enabling ParB-Smc association (**Supplementary Figure 3B**).

### Smc sequences crucial for ParB targeting

We next set out to identify Smc sequences responsible for association with ParB. Previous research uncovered a minimal Smc fragment that is proficient in *parS* targeting (Minnen *et al*., 2016). The fragment included the Smc head domain as well as about seventy amino acids of the head-proximal coiled-coil. *parS* targeting of this Smc fragment required the Walker B motif mutation E1118Q that prevents ATP hydrolysis but supports ATP-engagement of heads. Whether the head domain directly promotes ParB interaction or is merely required for ATP-mediated dimerization of the Smc fragment or for ATP-dependent DNA binding is however unclear (Vazquez Nunez *et al*., 2019).

Building on available structural information and prior experience with chimeric Smc proteins (Bürmann et al., 2017; Diebold-Durand et al., 2017), we constructed Smc chimeras with head-proximal sequences of *Bsu* origin and hinge-proximal sequences of *Spn* origin (**Figure 4A**). Junctions were chosen within the Smc joint or in its close proximity (ranging from Smc_234_-Smc_248_) with a crystal structure of the *Bsu* Smc joint (PDB: 5NMO) helping to keep amino- and carboxyterminal sequences in register (**Figure 4A, Supplementary Figure 4A**). All five chimeric proteins constructed in this fashion supported viability on nutrient-rich medium (**Supplementary Figure 4B**) implying that they are expressed and functional. This also highlights the lack of critical physical interactions between distal Smc parts (e.g. between the hinge and the head domains). Given the robust functioning of these chimeric proteins in the presence of *Bsu* ParB, we assessed their targeting directly by performing chromatin immuno-precipitation with antiserum raised against the ScpB protein followed by quantitative PCR (ChIP-qPCR), which revealed three patterns of distribution. Two chimeric proteins, Smc_205_ and Smc_234_, displayed reduced targeting to origin-proximal sites including the *parS_359_* site as well as at the *dnaA* and *dnaN* genes similar to the distribution found in a strain lacking *parB* (**Figure 4A**). These chimeric proteins thus appear to be unable to functionally interact with *Bsu* ParB. Two other chimeric proteins, Smc_241_ and Smc_248_, showed normal or near-normal distribution. The fifth construct, Smc_237_, displayed a *parS*-hyper-localization phenotype, suggesting defective protein release from *parS* sites. Taken together, these results show that sequences in the Smc joint and the immediately adjacent coiled coil mediate ParB-Smc interactions. The results are consistent with prior mapping studies based on non-functional and mutated Smc protein fragments (Minnen *et al*., 2016).

**Figure 4.**
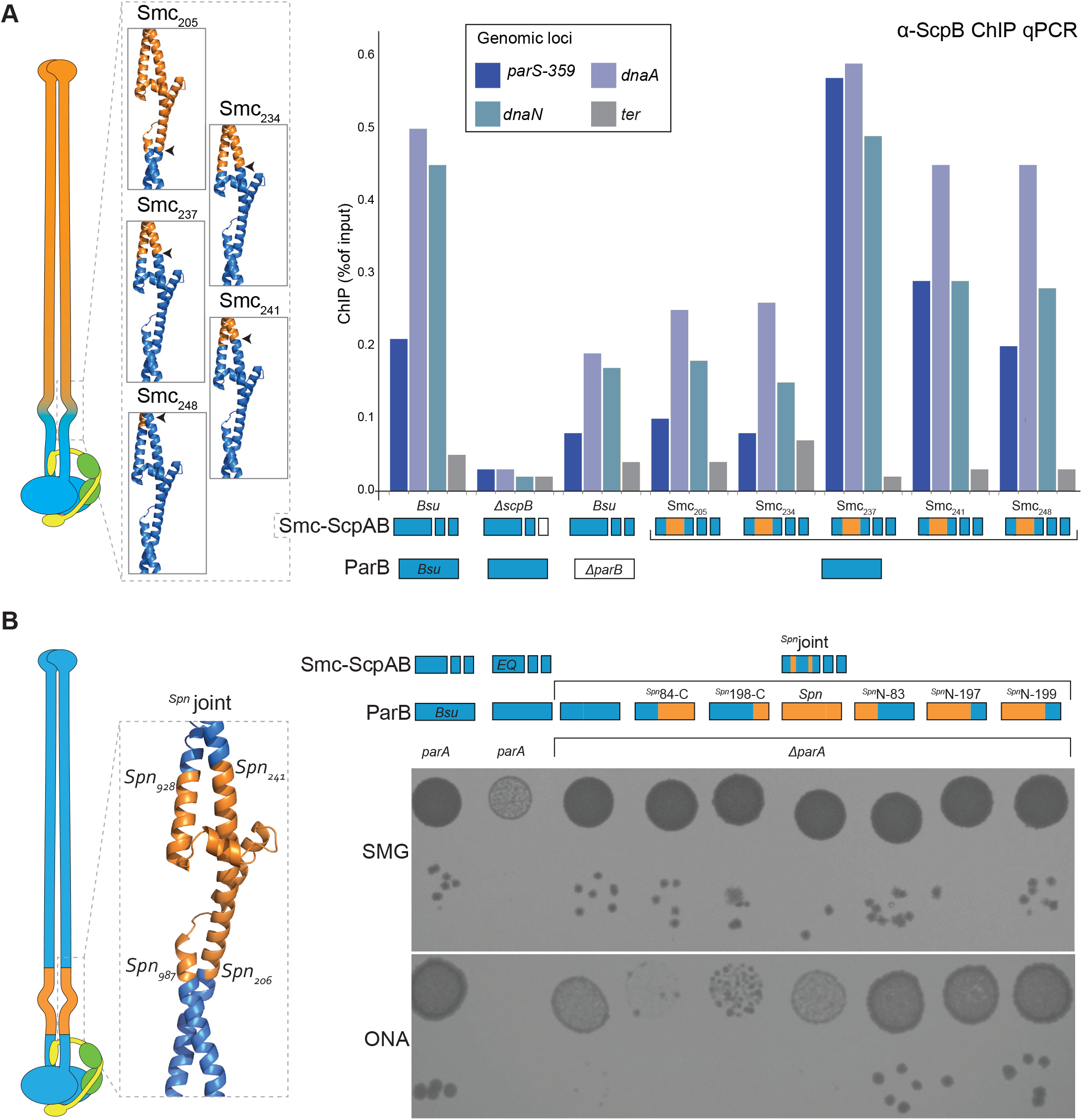
The Smc joint domain targets ParB. **A)** Left panel: Schematic (of Smc-ScpAB) and structure (of the *Bsu* Smc joint) denoting the construction of chimeric Smc proteins, blue colors indicating *Bsu* sequence identity, orange colors *Spn* origin. Right: Chromatin-immunoprecipitation coupled to quantitative PCR (ChIP-qPCR) using α-ScpB serum undertaken with chimeric Smc strains as denoted. **B)** Left panel: Schematic and structural model of Smc protein displayed as in A. Right: Viability assay by spotting of strains carrying ^*Spn*^joint in combination with indicated ParB chimeras. As in Figure 1B. For corresponding ChIP-qPCR results, see Supplementary Figure 4C.

### The Smc joint promotes ParB association

To test whether the Smc joint is sufficient to determine ParB specificity or whether head sequences are also necessary, we next constructed a chimeric Smc protein having only joint sequences of *Spn* origin (‘^*Spn*^joint’) (**Figure 4B**). Allelic replacement of *Bsu* Smc against this protein resulted in poor growth on nutrient-rich medium (**Figure 4B**) albeit noticeably better growth when compared to *Δsmc*.The ^*Spn*^joint protein is thus not fully functional which can likely be ascribed to combining multiple protein modifications (Smc_205_ and Smc_241_). Crucially, when combined with chimeric *parB* alleles having amino-terminal *Spn* sequences (^*Spn*^N-83, ^*Spn*^N-197 and ^*Spn*^N-199) it supported robust growth, while the converse *parB* alleles (^*Spn*^84-C, ^*Spn*^198-C and ^*Spn*^200-C) further decreased viability on nutrient-rich medium (**Figure 4B**). We found that these growth patterns correlated well with the chromosome distribution of ScpB in these strains as determined by ChIP-qPCR analysis (**Supplementary Figure 4C**). Of note, full-length *Spn* ParB did not significantly improve the viability of the ^*Spn*^joint strain or the recruitment of ^*Spn*^Smc-ScpAB, possibly indicating that C-terminal ParB sequences might contact Smc-ScpAB sequences outside the Smc joint and thus contribute to Smc-ParB associations (see below). Together, the above experiments show that sequences in the Smc joint and the ParB CTP-binding domain need to be matched to enable productive ParB-Smc contacts. This demonstrates that a direct physical interaction between these regions is necessary for optimal function. The head domains likely contribute to the targeting of minimal Smc fragments indirectly, by mediating Smc dimerization and DNA binding (Vazquez Nunez *et al*., 2019)

### Proximity of ParB and Smc detected by chemical cross-linking

Finally, we applied *in vivo* cross-linking to complement the genetics and to further fine-map the Smc-ParB interface. Given the lack of structural information on the interface, we approached the problem in two steps. We first cross-linked candidate cysteine residues in ParB to lysine residues in Smc using the heterobifunctional Lys-Cys cross-linker SMCC (**Figure 5A**). In a second step we used the homobifunctional Cys-Cys cross-linker BMOE (**Figure 5D**) on combinations of ParB(Cys) and Smc(Cys) mutants. To enhance the fraction of cellular Smc proteins localized at *parS*, (Minnen *et al*., 2016), we utilized an EQ mutant also defective in hinge dimer formation (‘mH’) (Hirano and Hirano, 2002). The Smc protein also harbored a HaloTag (‘HT’) for quantitative detection of cross-linked species. Seventeen residues on the surface of ParB were selected to evenly cover the N-terminal sequence of ParB (**Figure 5B**). Only when cross-linking with ParB(A29C), ParB(K67C), and ParB(S100C), we observed an additional, more slowly migrating species of Smc(mH/EQ)-HT (**Figure 5B**). We note that these three residues happen to localize to patches 1, 3, and 4, respectively (**Figure 3A**). They are closely juxtaposed to each other on the surface of ParB (**Figure 5C**) and presumably also located in proximity of one or more lysine residue(s) of Smc.

**Figure 5.**
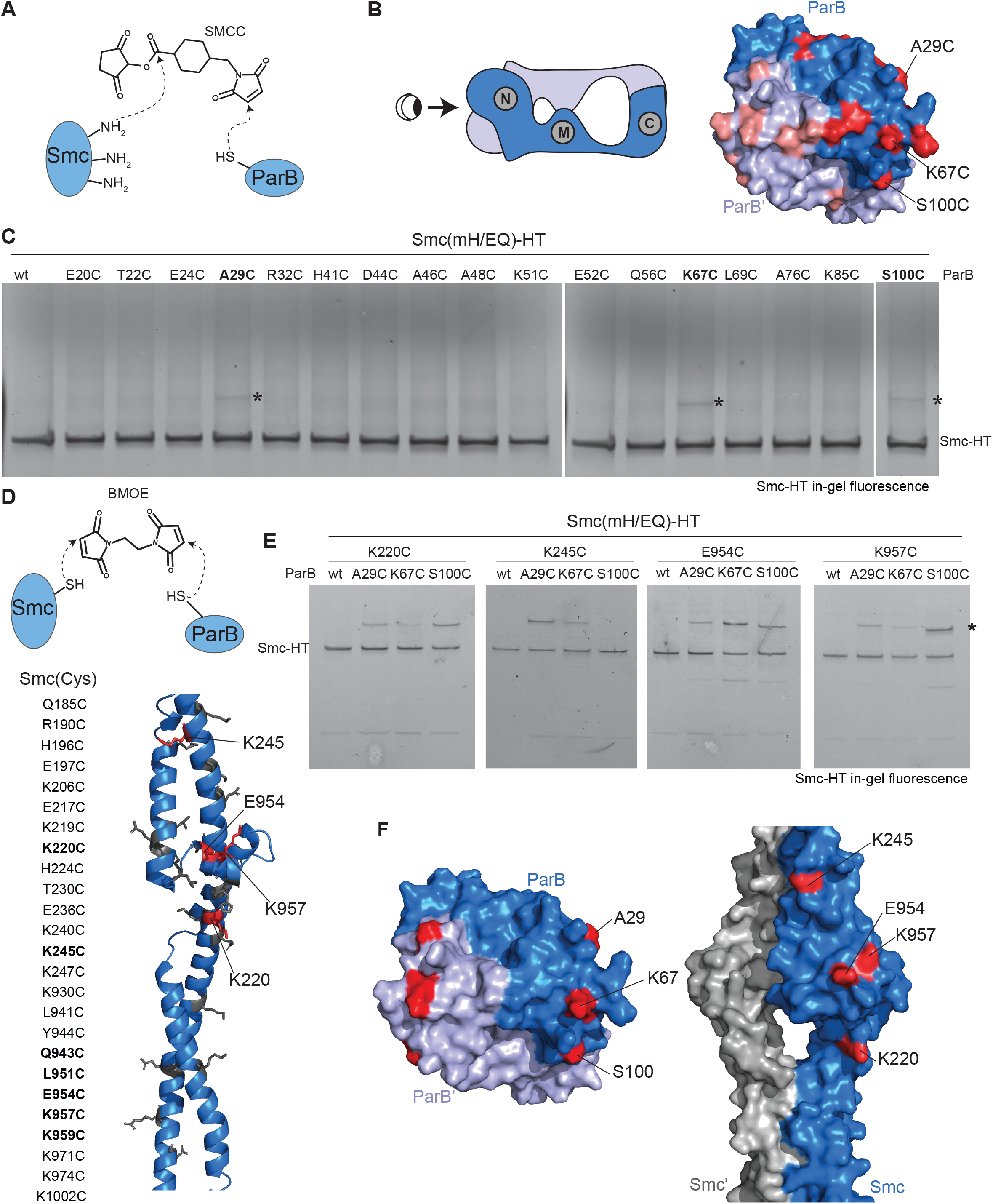
*In vivo* cross-linking supports the interaction interfaces found by genetics. **A)** Schematic of chemical cross-linking by the heterobifunctional molecule SMCC. **B)** Candidate ParB cysteine residues and their position (in red colors) on the ParB-CDP dimer (with chains in blue and grey colors in surface representation). **C)** SMCC cross-linking using ParB(Cys) mutants as indicated and detected by in-gel fluorescence detection of Smc(mH/EQ)-HT (‘Smc-HT’) protein. Higher molecular weight species appearing upon cross-linking are indicated by asterisks. **D)** Schematic of BMOE cross-linking chemistry (top panel) and candidate Smc(Cys) residues and their distribution on the Smc joint structure (in cartoon representation). **E)** BMOE cross-linking using combinations of ParB(Cys) and Smc(Cys) mutants as indicated. Samples were enriched for ParB interacting material by incubation with α-ParB antibody coupled Dynabeads. Detection by in-gel fluorescence of Smc(mH/EQ)-HT (‘Smc-HT’) protein. Cross-linked ParB-Smc species are indicated by asterisks. **F)** Positioning of identified cross-linking residues on ParB (left panel, as in B) and Smc (right panel, Smc joint in the rod configuration in surface representation).

To identify the Smc residues that are in proximity of ParB, we selected twenty-five candidate residues located on both α-helices of the joint to be mutated to cysteines (**Figure 5E**). The mutations were generated in the Smc(mH/EQ)-HT protein and combined with the three ParB(Cys) mutants identified above (**Figure 5F**). To improve the detection of cross-linked products, we enriched for ParB species by pull-down assays with serum raised against the ParB protein. Comparable results were however obtained without enrichment (**Supplementary Figure 5B, 5C**). We found comparatively robust cross-linking with only four Smc(Cys) mutations: K220, K245, K954, and K957 (**Figure 5F, Supplementary Figure 5B, 5C**). These four residues were broadly distributed along the Smc joint and present in both coiled coil α-helices. However, all four were exposed on the same side of the Smc joint surface (**Figure 5F**).

### Predicting the structure of the joint-ParB interface

We next applied structure prediction to the minimal interacting sequences as identified by our mapping experiments (Figure 6A). Structure interface prediction became feasible recently with the advent of AlphaFold-Multimer (AF-Multimer) (Evans et al., 2021). The folds of the two individual chains of the ParB-Smc structure are predicted with high confidence and the predictions superimposed well with published crystal structures of the Smc joint (PDB: 5NMO) and ParB NM (PDB: 6SDK), respectively (Supplementary Figure 6B, C). AF-Multimer consistently predicted a tightly fitted heterodimer of the two protein fragments, albeit with a lower level of confidence for the interface prediction (Supplementary Figure 6A). Consistent with our mapping experiments, the ‘joint-ParB’ interface is formed by the ParB N domain and the middle and head-distal regions of the Smc joint. The C_α_-C_α_ distances measured for four efficiently cross-linked cysteine pairs (Supplementary Figure 5B) (^~^8 to 17 Å) fit well in the range for BMOE cross-linking (Figure 6B) (Diebold-Durand *et al*., 2017). Moreover, ParB residues identified through the matching of sequences in chimeric proteins are found directly at the interface (Figure 6C). The AF-Multimer model thus likely closely resembles the ParB-Smc structure formed during the recruitment of Smc-ScpAB to the ParB partition complex.

**Figure 6.**
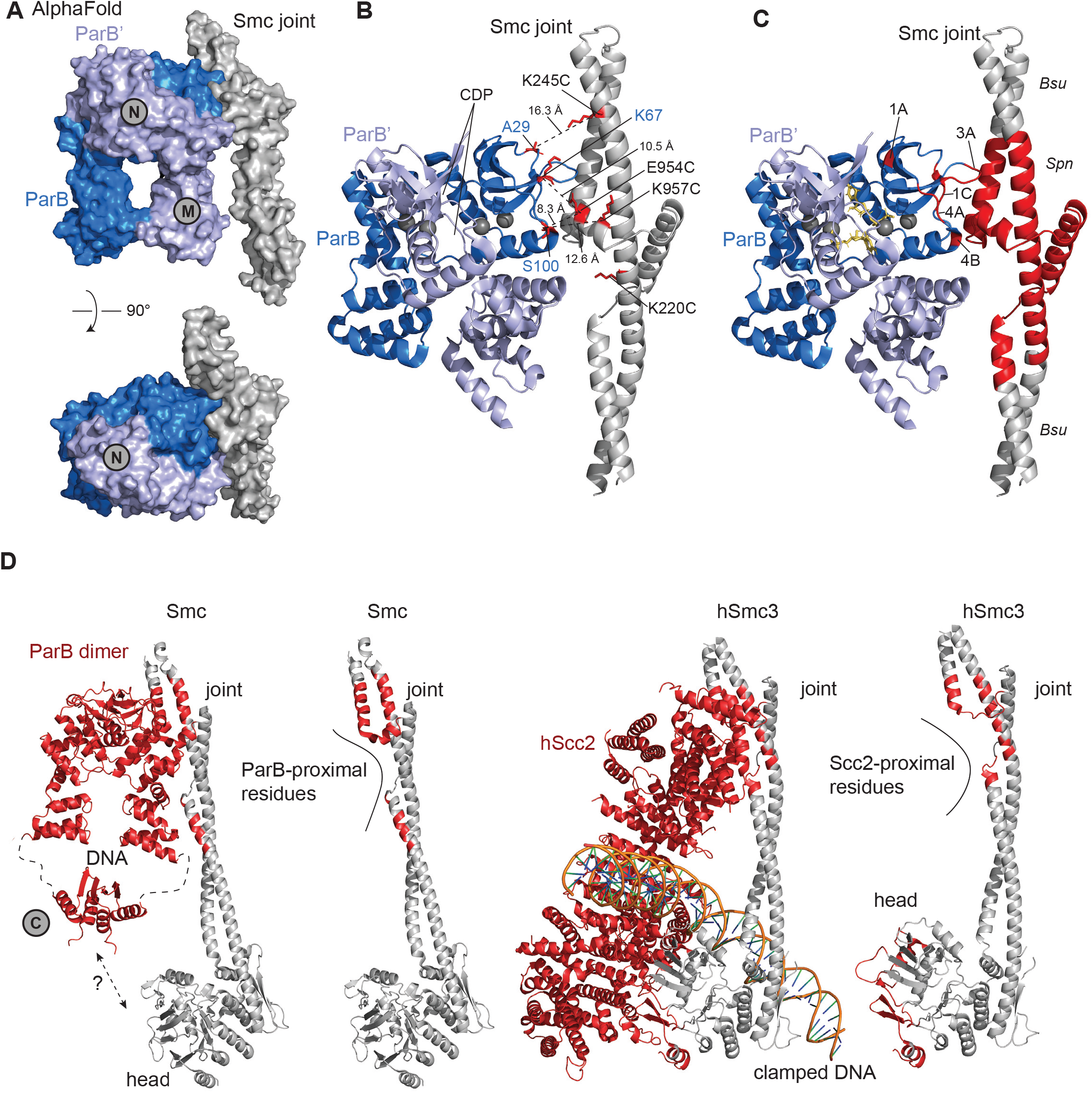
Structure prediction and comparison. **A)** A reconstruction of an Smc-ParB sub-complex obtained by superimposition of a crystal structure of the ParB NM domain dimer (PDB: 6SDK) with a joint-ParB heterodimer predicted by AF-Multimer (see Supplementary Figure 6A) in surface representation in side view (top panel) and top view (bottom panel). The Smc chain is displayed in grey colors, the ParB chains in dark and light blue colours, respectively. **B)** The Smc-ParB sub-complex shown in cartoon representation with residues used for cysteine cross-linking experiments displayed as sticks. C_α_-C_α_ distances (in Å) are indicated by dashed lines. **C)** Same as in panel B with Smc and ParB residues identified by genetic sequence matching displayed in red colors. **D)** Side-by-side comparison of the joint-ParB interface (left panels) and the joint-Scc2 interface in human cohesin (right panels). ParB- and Scc2-proximal residues on the Smc and Smc3 joint (C_α_-C_α_ distance < 10 Å), respectively, are indicated in red colors. A Scc2/DNA sub-structure of ATP-engaged human cohesin is displayed (PDB: 6YUF) (right panel). Only the Smc3 and Scc2 subunits are shown for simplicity. For direct comparison, the SMC subunits are also displayed in isolation.

## Discussion

Revealing how SMC complexes load onto DNA is vital for a basic understanding of the mechanism of DNA loop extrusion. Here we provide first insights into how a *bona fide* DNA loading factor (*i.e*. ParB) delivers DNA to an SMC complex for the initiation of DNA loop extrusion. We identify the joint-ParB interface as a major determinant for Smc targeting in bacteria.

### DNA loading by Smc-ScpAB

A key question on the mechanism of DNA loop extrusion is how chromosomal DNA arrives at the DNA clamping site on top of ATP-engaged head domains (Vazquez Nunez *et al*., 2019). There are two possible scenarios (irrespective of SMC-DNA topology) (**Figure 7A**): A DNA double helix is transferred between disengaged heads and then becomes clamped by ATP-engaging Smc heads (scenario ‘1’ in Figure 7B). Alternatively, heads first ATP-engage with one another, thus closing the route between the heads. A loop of DNA then engages directly from the coiled-coil proximal side (scenario ‘2’ in Figure 7B) (Vazquez Nunez *et al*., 2019). Both scenarios are in principle compatible with the joint-ParB interface. Passage of DNA between the heads (scenario 1) has been suggested for yeast cohesin based on a reconstituted DNA loading reaction using purified components (Collier et al., 2020). However, it is not obvious how the joint-ParB interface would promote or even direct such a passage mechanism because formation of joint-ParB interface prior to DNA clamping would position DNA across the heads rather than between them. Also, it is unclear how this would lead to entrapment of chromosomal DNA by the Smc-ScpA ring (Vazquez Nunez *et al*., 2019). The entry of DNA at the top of the heads (scenario 2) has been suggested for DNA-end recognition by Rad50-Mre11 (Käshammer et al., 2019). At a DNA double strand break, the DNA end can thread into the interarm space. In case of Smc-ScpAB, however, a pre-formed DNA bend or loop has to thread into the interarm space. It is conceivable that such loops readily form in the partition complex. In case of phage P1, DNA between *parS* site motifs is bent significantly at a IHF protein binding site (Surtees and Funnell, 2001). Moreover, trans-contacts between ParB clamps may form or stabilize such loops. The joint-ParB interface seems to be ideally positioned to guide a DNA loop into an opened SMC compartment, although the details of the DNA passage remain largely unclear.

**Figure 7.**
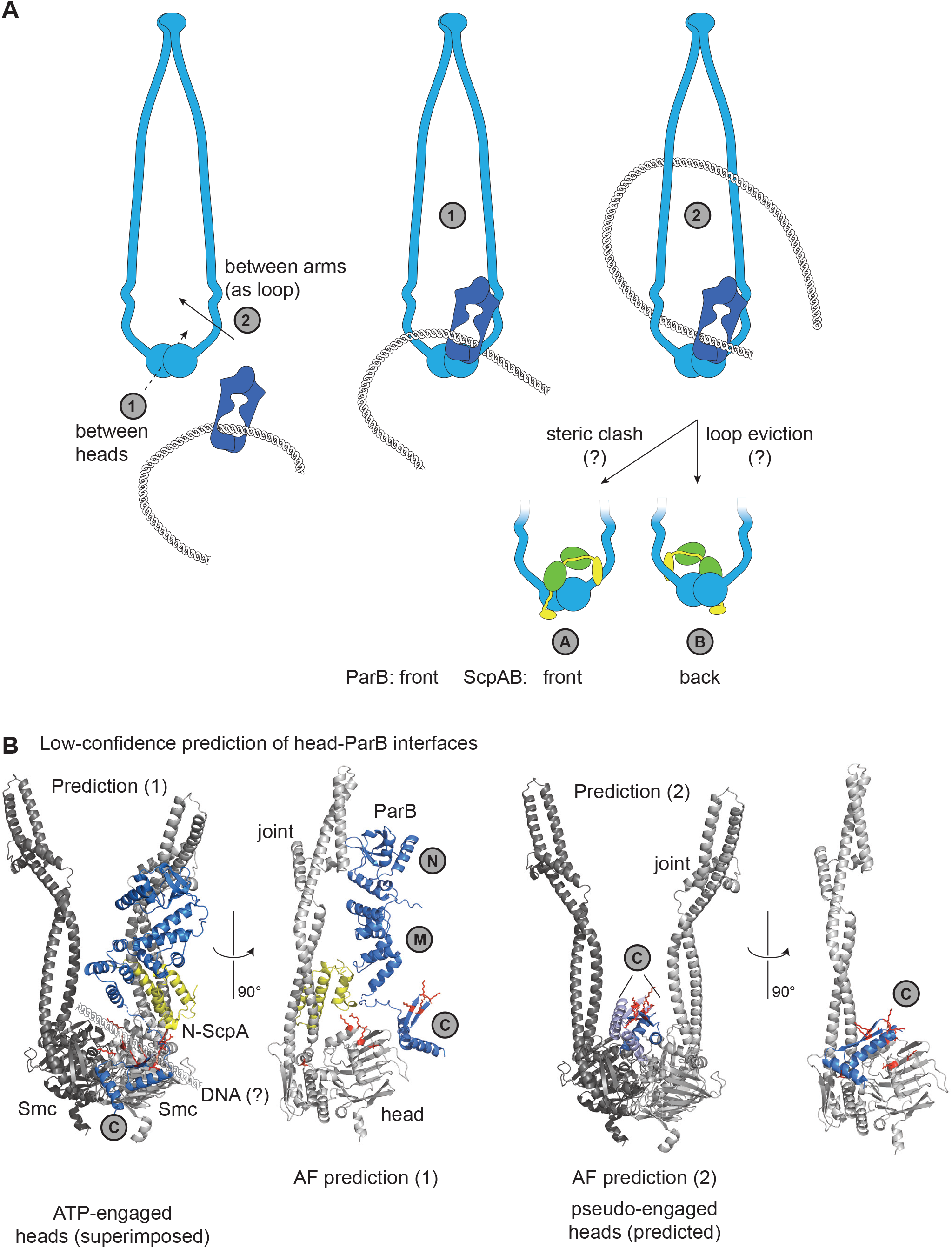
Models and low-confidence AF predictions. **A)** Putative models for the contact between ParB-clamped DNA and the Smc dimer. For simplicity, ScpAB is omitted from some representations and indicated separately in the bottom panels. Two scenarios are considered: DNA passage between disengaged heads (‘1’) and insertion of a DNA loop into the Smc interarm space (‘2’). The products of these reactions are shown in the middle and right panels. ParB is shown to interact with the right Smc (ParB in ‘front’ of Smc). ScpAB can either be associated on the same side (‘front’) or other side (‘back’). Possible variations of these scenarios with pseudo-topological and non-topological modes of association between Smc-ScpAB and DNA are not shown for the sake of simplicity. B) AF-Multimer predictions of head-ParB interfaces: input sequences for prediction (1): Smc-head-joint, full-length ParB and the N-terminal domain of ScpA (left panels); for orientation a second Smc-head-joint protein is superimposed in an ATP-engaged arrangement (left panel); the putative location of DNA is indicated (left panel). Input sequences for prediction (2): Smc-head-joint as dimer with the ParB C domains as dimer. Note that the Smc dimer is predicted in a pseudo-engaged state (despite the absence of ATP).

The symmetric nature of the bacterial Smc dimer (**Figure 7A**) allows for two joint-ParB interfaces on the asymmetric Smc-ScpAB holo-complex. ParB may either bind on the same side as ScpAB (‘front’), which would likely result in a steric clash since the middle part of ScpAB occupies a similar area on top of ATP-engaged Smc heads (**Figure 7A**) (according to the position of the corresponding kite subunits in DNA-clamping MukBEF) (Bürmann et al., 2021). Alternatively, ParB may approach from the other side (‘back’) and thus avoid a steric clash. However, in this scenario, subsequent DNA translocation (without prior conversion to a topological DNA-Smc association – ‘loading’) would evict the newly captured DNA loop from the Smc complex and thus be counterproductive at least according to the DNA-segment-capture model (Diebold-Durand *et al*., 2017; Marko et al., 2019; Nomidis et al., 2021). We thus propose that ParB substitutes for ScpAB in DNA clamping during chromosomal loading, possibly analogous to Pds5 substituting for Scc2 in cohesin unloading from DNA (Wells et al., 2017). We previously found that ScpA is not required for *parS*-targeting of the Smc(EQ-mH) protein (Minnen *et al*., 2016). This option is also supported by experiments with engineered asymmetric Smc dimers which harbor only the joint in the ν-Smc protein substituted for *Spn* sequences (**Supplementary Figure 7A**) (Bürmann *et al*., 2013). However, the results are not fully conclusive because the joint substitution in the κ-Smc also performs well with one particular chimeric ParB allele, probably indicating that multiple ParB-Smc contacts or multiple ParB-Smc states together support loading (**Supplementary Figure 7B**).

The inherent symmetry of the Smc and ParB dimers may also allow for the simultaneous engagement of two joint-ParB interfaces, thus possibly stabilizing the Smc-ParB association. Under the reasonable assumption that the ParB N-M architecture is more or less rigid (Soh *et al*., 2019), this is only possible after major reorganization of the Smc coiled coils requiring an X-shaped arrangement of the Smc proteins in the dimer (**Supplementary Figure 7C**). Whether such an extreme coiled coil configuration occurs even transiently is doubtful. Another possibility is that a single ParB clamp recruits two Smc complexes *e.g*. to build a dimeric motor complex for bidirectional translocation. This however also seems unlikely considering the high local concentration of ParB dimers that would compete for Smc binding. More likely, the ParB dimer may be handed over from one Smc arm the other as the ParB clamp threads through the Smc dimer during DNA loading.

### Key functions of the SMC joint in DNA recruitment, loading, translocation and unloading

The SMC joint serves as a key binding platform also in other SMC complexes. MatP protein (bound to *matS* sites) is an unloading factor for MukBEF, which releases it from the chromosome in the replication terminus region (Lioy et al., 2018). The AcpP protein has an important stimulatory effect on the ATPase activity of MukBEF (Josh et al., 2021). Both proteins bind to the joint in MukBEF (the two joints actually) (Bürmann *et al*., 2021). The hawk subunit Scc2 is a loading as well as processivity factor for DNA loop extrusion by cohesin (Davidson et al., 2019). It forms an interface with the Smc3 joint in the DNA-clamping, ATP-engaged state of cohesin (Higashi et al., 2020; Shi et al., 2020). An equivalent joint-hawk interface is also found in condensin (Lee et al., 2021; Shaltiel et al., 2021). To understand how closely these joint interfaces might be related to one another, we superimposed the joint domain in these structures. Superimposition with MukB (PDB: 7NZ3) is challenging due to the significantly diverged structure of the joint (Bürmann *et al*., 2021). The Smc3 joint (PDB: 6YUF) superimposed significantly better with the Smc joint and intriguingly showed that the hawk-joint interface overlaps well with the joint-ParB interface (**Figure 6D**) (Higashi *et al*., 2020). SMC loading (in cohesin and Smc-ScpAB), translocation (in cohesin and condensin) as well as SMC unloading (in MukBEF) are thus linked to related joint interfaces, highlighting the importance of the joint for SMC DNA transactions. While ParB and hawk proteins are structurally and phylogenetically unrelated, they use equivalent binding sites on SMC. The binding of co-factors to the joint for DNA clamping might thus be a general feature of many or all SMC complexes.

### Additional ParB-Smc interfaces?

To identify putative additional contacts between ParB and Smc-ScpAB that may help to guide DNA into the Smc complex, we ran structure predictions with larger input sequences. In some of the AF-Multimer predictions, we found the ParB C domain docking onto the side of the Smc heads (via a ‘head-ParB interface’) (**Figure 7B**, left panels). Intriguingly, in this scenario, the DNA binding surface of ParB C domain aligns side by side with the DNA binding surface on the Smc heads, implying that the proteins form a composite DNA binding surface. Other predictions with two chains of Smc (heads only) and ParB showed a dimer of ParB C domains on top of (pseudo-engaged) Smc heads in a position normally occupied by clamped DNA (**Figure 7B**, right panel). Together, these predictions indicate that the flexible nature of the M-to-C connections in the ParB dimer allows for the ParB-clamped DNA to be located at the Smc heads, either being also clamped by Smc heads or held in place by additional ParB-Smc contacts. Conceivably, multiple ParB-Smc contacts are formed sequentially as DNA is being threaded into the Smc compartment.

### The ParB CTP binding domain

The N-terminal CTP-binding domain of ParB has a crucial role in the targeting of Smc to the chromosome. It is located farthest away from the clamped DNA, thus possibly reaching out from the partition complex to contact and capture free Smc dimers (**Figure 7A**). In close vicinity lies the unstructured N-terminal peptide of ParB that stimulates ParA ATP hydrolysis, potentially suggesting a mutually exclusive binding of ParA and Smc to ParB and antagonistic regulation. Notably, ParA chromosome segregation requires ParB CTP hydrolysis while Smc recruitment does not, indicating further opportunities for antagonistic regulation (Antar *et al*., 2021).

### A minimal system for chromosome folding - dispensability of host factors

We found that the modules for sister chromosome individualization are remarkably robust and able to segregate chromosomes in a distantly related host bacterium, implying that direct binding of host factors is dispensable for the essential function. Moreover, such host factors do not interfere with the basal activity of ParB and Smc-ScpAB. DNA loop extrusion by Smc-ScpAB thus occurs at least partly unhindered by orthologous obstacles on the chromosome. This implies that overcoming such obstacles does not require dedicated bypass mechanisms with physical contacts between DNA motor and obstacle (Anchimiuk *et al*., 2021; Brandão et al., 2021). Bypassing of DNA binding factors while forming chromosomal loops thus appears to be an inherent propensity of Smc-ScpAB.

### Mapping weak binding interfaces by gene transplantation and structure prediction

How the different players in cellular pathways interact with one another to support optimal coordination of their activities is often poorly understood. We initially aimed to detect the Smc-ParB interaction by performing biophysical interaction studies (including pull-down, co-elution, anisotropy, Bio-layer interferometry) using purified components and cofactors. However, all of our attempts were unsuccessful, possibly owing to the weak and very transient nature of the association or the dependence on a cofactor or a posttranslational modification. ParB self-concentrates to unusually high cellular concentrations within the partition complex (estimated to be as high as 10 mM at least for a plasmid ParB protein) (Guilhas et al., 2020), thus possibly bypassing the need for a high affinity contact. Considering this high local concentration of ParB, it is conceivable that the dissociation constant (K_D_) value for such an interaction is in the high μM or even mM range. A more stable interaction of ParB and Smc would have limited positive impact upon Smc recruitment but may hinder or even block its subsequent release. Similar conditions are likely found in proteins undergoing liquid-liquid phase separation. Our approach based on gene transplantation and cross-linking may be more widely applicable to study the recruitment of factors by biological condensates formed by liquid-liquid phase separation (Feng et al., 2019). When combined with structure prediction, this approach may be particularly powerful.

## Acknowledgements

We are grateful to Frank Bürmann for comments on the manuscript and all members of the Gruber lab for stimulating discussions and feedback. We thank the Jan-Willem Veening lab for help with imaging. This work was supported by the Swiss National Science Foundation (197770 to S.G.) and the European Research Council (724482 to S.G.).

## Author contributions

FB cloned the recombinant DNA constructs, obtained the genetically modified *B. subtilis* strains, and performed experiments except for 3C-Seq. AA implemented 3C-Seq experiments and analysis. MLDD established in vivo Lys-Cys crosslinking with SMCC. FB and SG prepared the figures and wrote the manuscript text with input from all authors. SG supervised the work and obtained funding.

## Data Availability

Raw sequencing data obtained in this work can be found on NCBI-SRA (accession number: GSE190491). All other raw data is available via Mendeley (DOI: 10.17632/3k5sffj2w4.1).

## Supplementary Figure legends

**Supplementary Figure 1.**
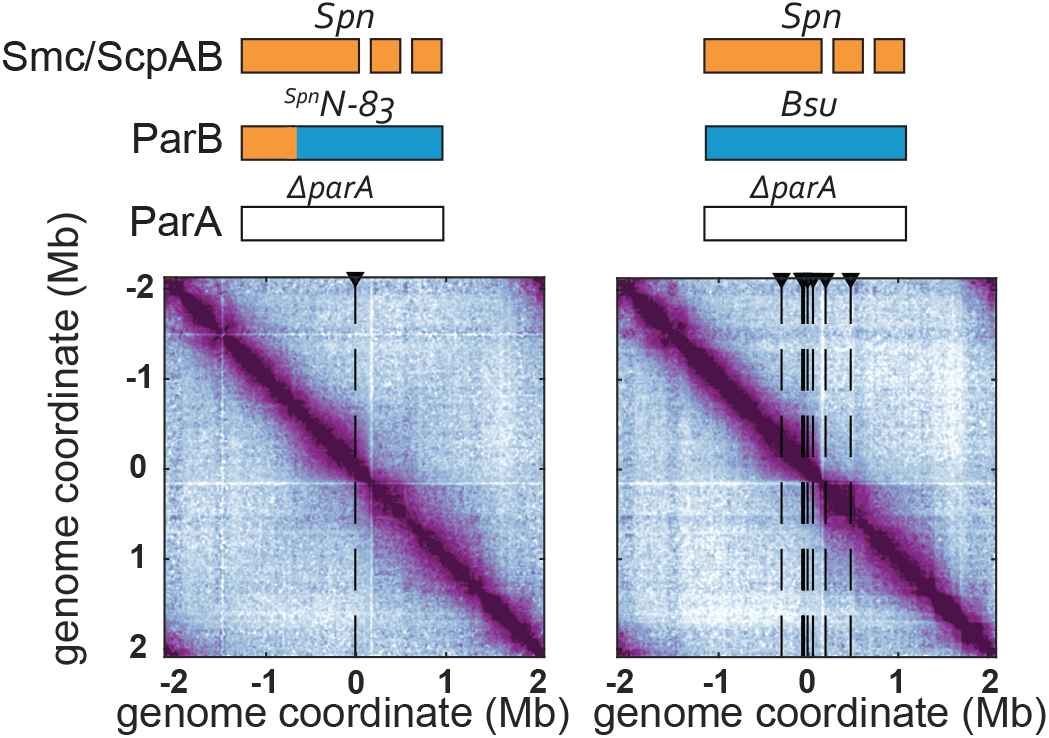
Normalized 3C-seq contact maps of strains with indicated genotypes grown exponentially as in Figure 1C.

**Supplementary Figure 2.**
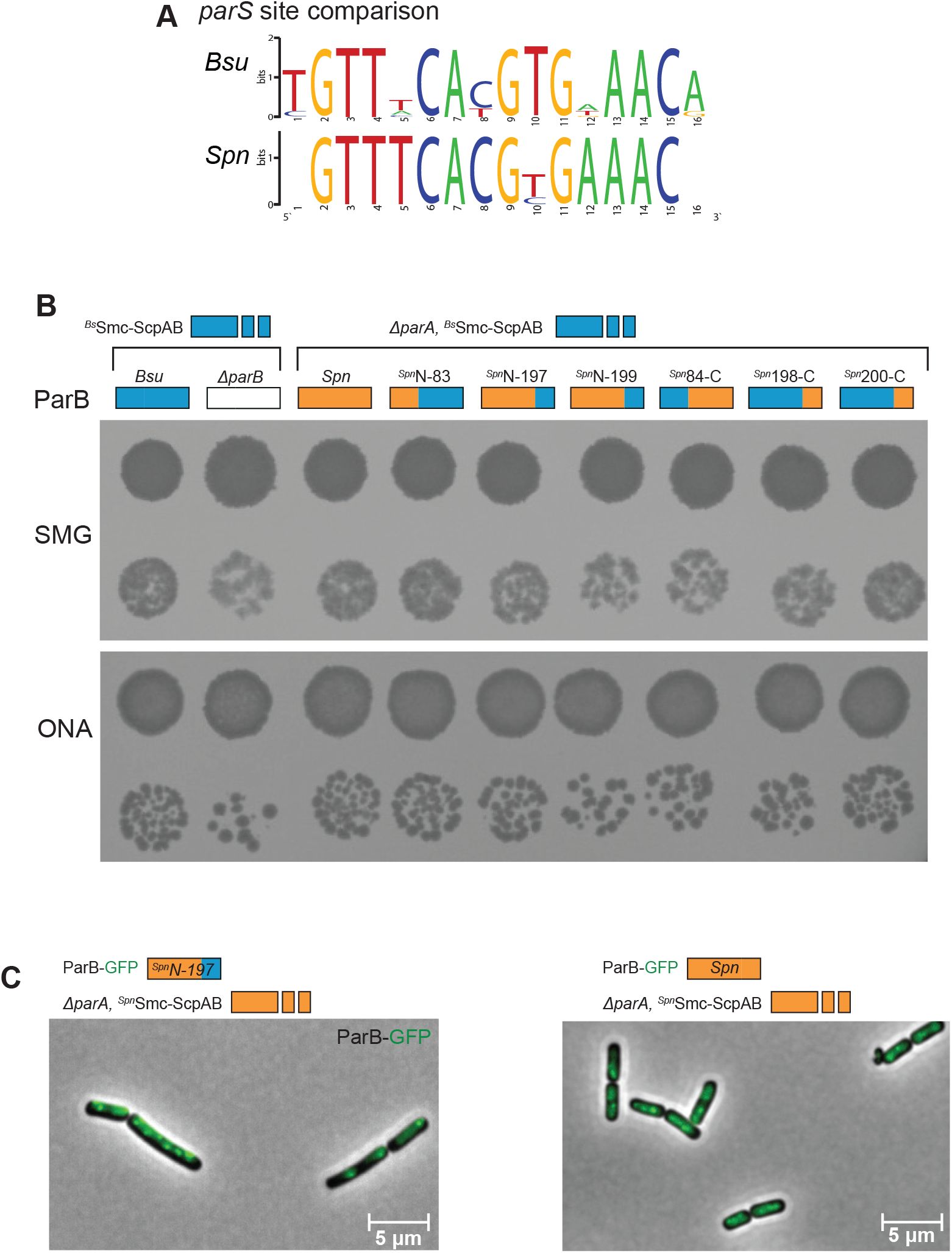
GFP-tagged chimeric constructs are expressed in *B. subtilis*. **A)** Sequence logo for the alignment of *Bsu* and *Spn parS* sites, respectively. **B)** Viability of cells harboring different *Bsu/Spn* chimeric ParB sequences. As in Figure 1B. **C)** Microscopy image *B. subtilis* cells containing the ^*Spn*^Smc-ScpAB with either ^Spn^N-197 or ^*Spn*^ParB proteins fused to GFP. As in Figure 2C.

**Supplementary Figure 3.**
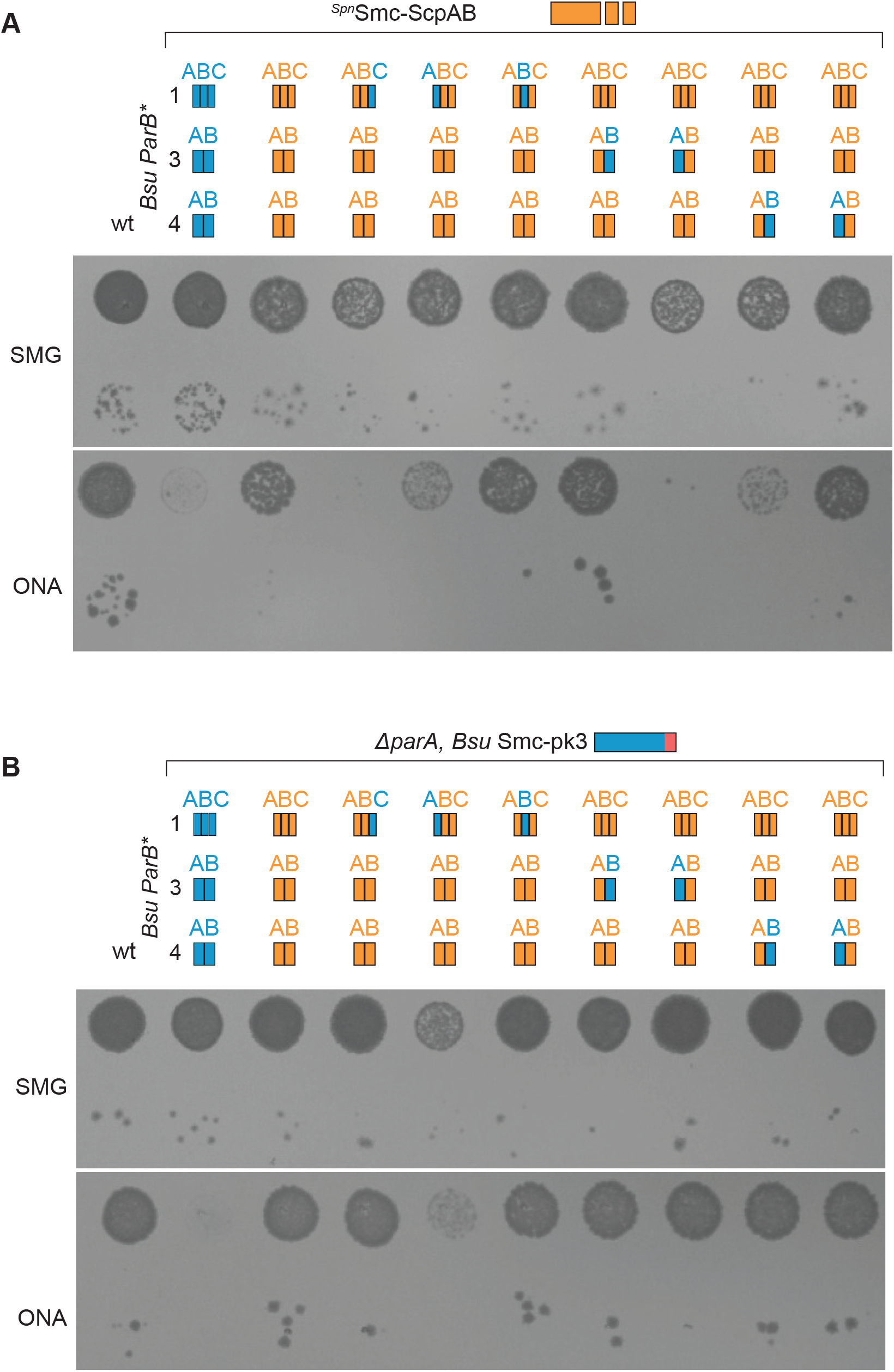
Mapping of ParB residues involved in Smc interaction. **A)** Spotting assay of strains carrying ^*Spn*^Smc-ScpAB, ParB chimeras as well as an intact *parA* genes. As in Figure 3C but with ParA protein being present. **B)** Spotting assay of strains carrying the ^*Bsu*^Smc-pk3 allele as well as *parA* deletion and ParB chimeras as indicated.

**Supplementary Figure 4.**
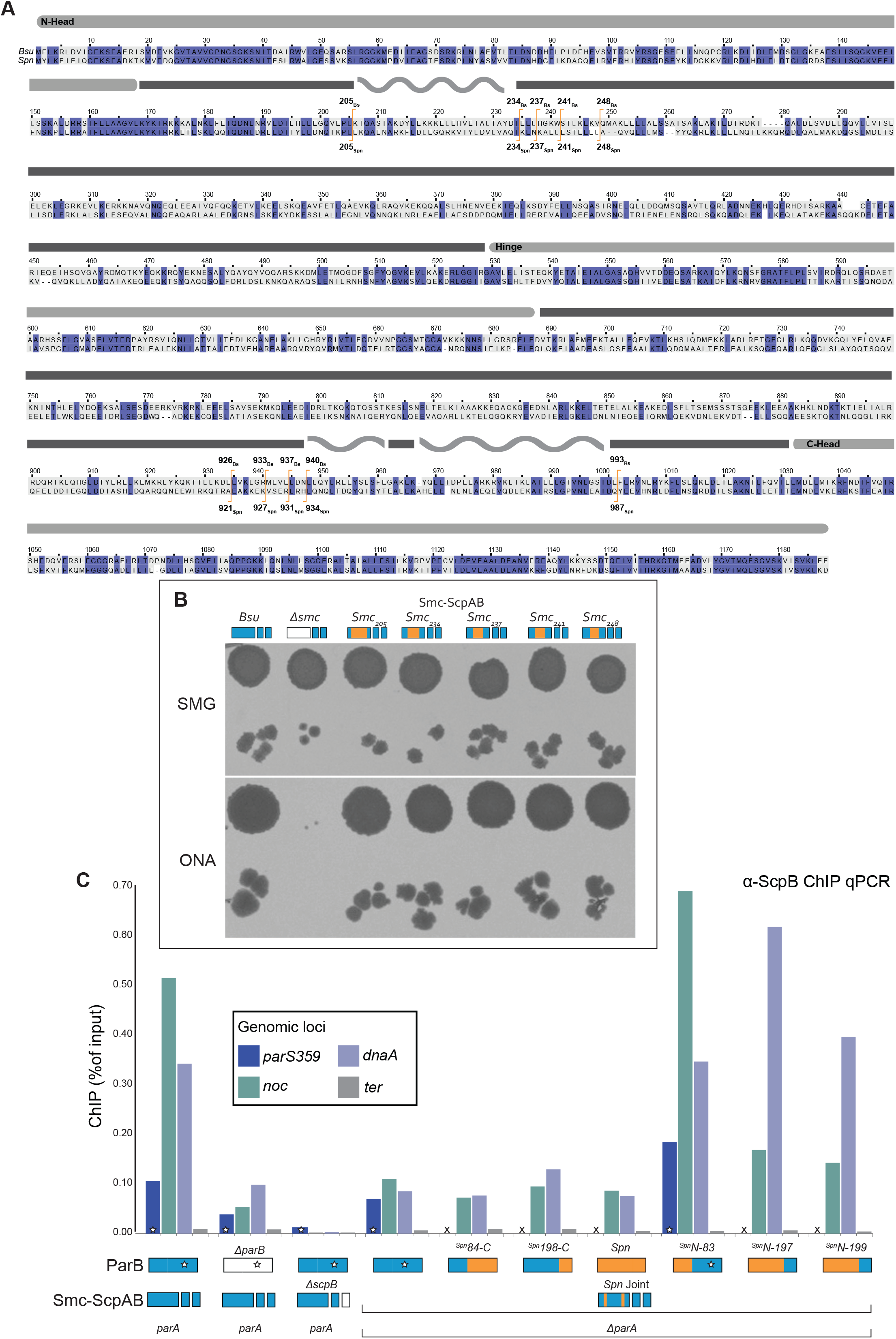
Construction and analysis of chimeric Smc proteins. **A)** Alignment of *Bsu* and *Spn* Smc amino acid sequences. Identical residues are highlighted by blue background. Joint domain is indicated above. Transitional points between the protein sequences are indicated in orange. **B)** Viability of strains with Smc chimeras and wt *Bsu* ParB. **C)** Chromatin-immunoprecipitation coupled to quantitative PCR (ChIP-qPCR) using α-ScpB serum undertaken on ^*Spn*^joint strains containing ParB chimera proteins as indicated. Strains identical to the ones used for spotting in Figure 4B.

**Supplementary Figure 5.**
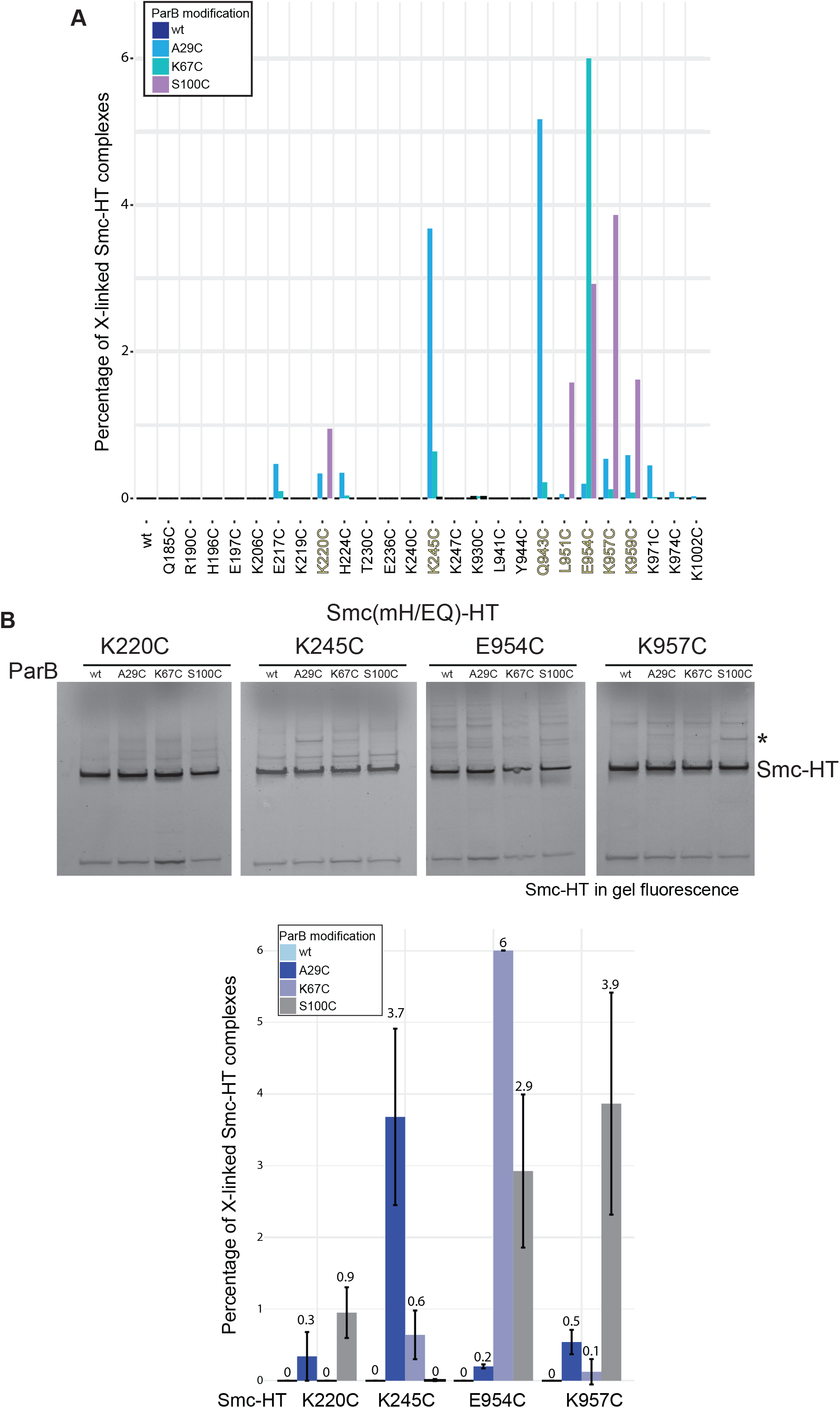
Detection of Smc-ParB cross-linking. **A)** Screening combinations of ParB(Cys) and Smc(Cys) residues for BMOE cross-linking based on the detection of Smc(mH/EQ)-HT (‘Smc-HT’) protein by in-gel fluorescence. **B)** Cross-linking of selected Smc(Cys)-ParB(Cys) combinations as in Figure 5E but without pre-enrichment by ParB immunoprecipiation. Detection by in-gel fluorescence of Smc(mH/EQ)-HT protein. Representative gel images are shown (top panel). Quantification of BMOE cross-linking (bottom panel).

**Supplementary Figure 6.**
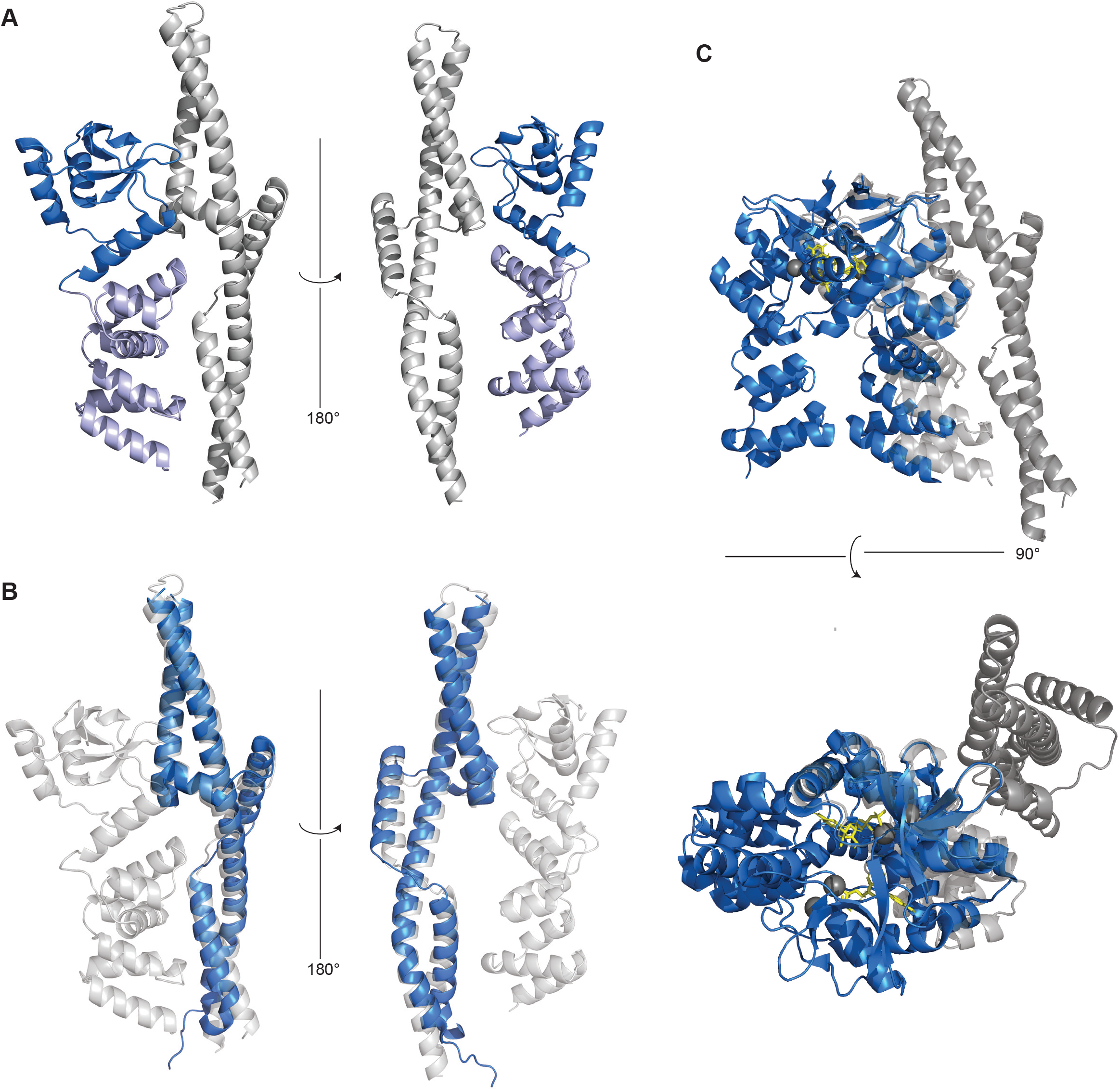
Structure prediction and superimposition. **A)** A representative model of the Smc joint complex with the ParB NM domain fragment obtained by AF-Multimer in cartoon representation front and back views (left and right panel, respectively). The Smc chain is displayed in grey colors, the ParB N and M domains in dark and light blue colors, respectively. **B)** Superimposition of the model shown in A (in grey colors) with the crystal structure of the Smc joint (PDB: 5NMO) (in blue colors). **C)** Superimposition of the model shown in A (in grey colors) with the crystal structure of the ParB NM domain dimer (PDB: 6SDK) (in blue colors).

**Supplementary Figure 7.**
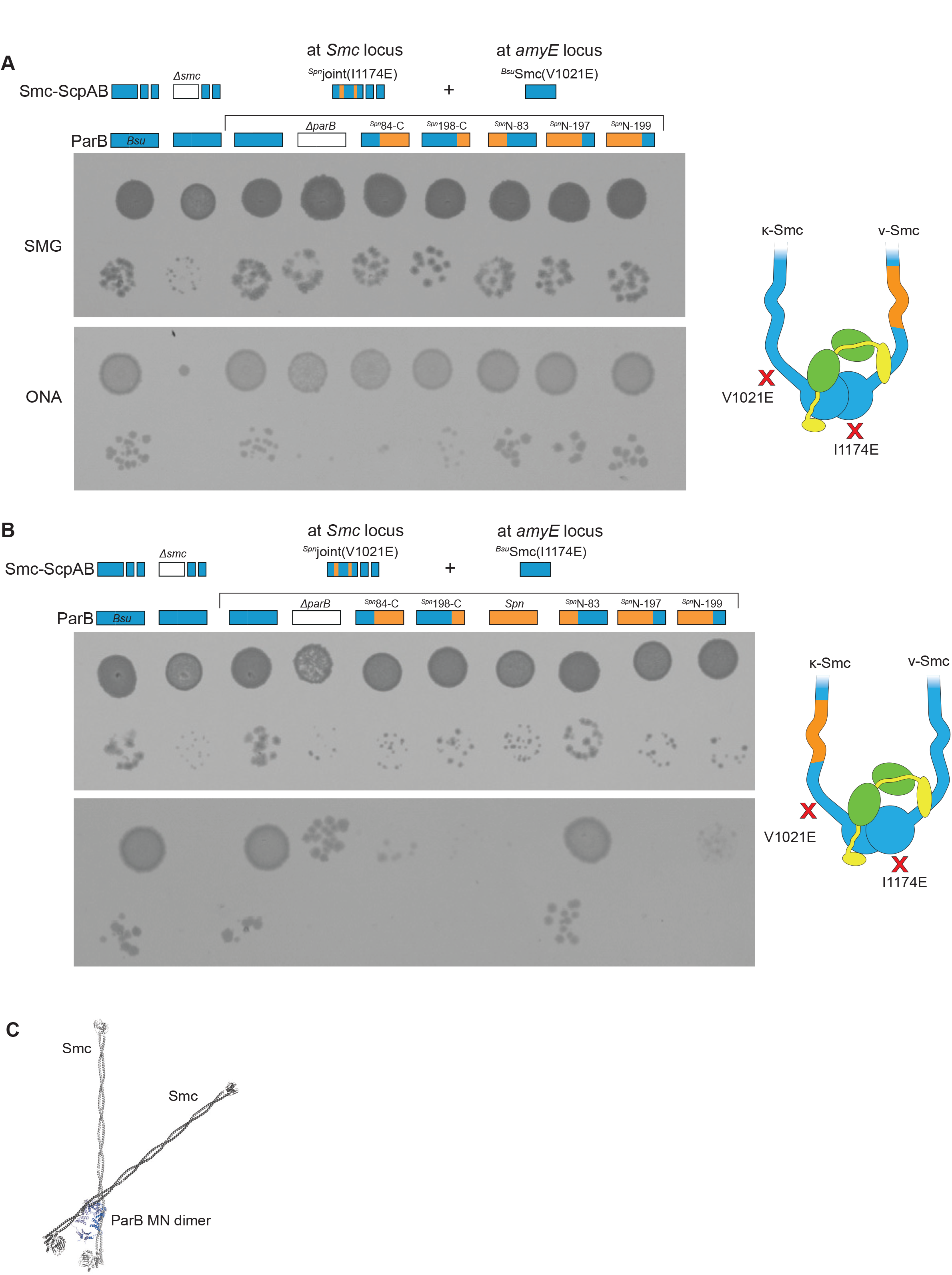
Mapping of ParB residues involved in κ-Smc and η-Smc interactions. **A)** Spotting assay of strains carrying two different Smc alleles: an ^*Spn*^Joint gene harboring the I1174E mutations expressed from the endogenous locus and an ectopic copy of the *Bsu* Smc gene carrying the V1021E mutation. The V1021E and I1174E point mutations block N-ScpA and C-ScpA binding, respectively. Only heterodimeric Smc dimers assembly functional Smc-ScpAB complexes. ParB chimeras as indicated. **B)** As in A but with V1021E and I1174E mutations swapped. **C)** Reconstruction of a ParB-Smc 2:2 complex using two Smc monomers (rod state) (in light and dark grey colors, respectively) and the ParB NM crystal structure (PDB: 6SDK) (in light and dark blue colors).

## Materials and Methods

### Strain construction

*B. subtilis* strains utilized in this work originate from the 1A700 isolate. Natural competence was used to engineer strains at the *smc*, *scpAB, parAB* and *amyE* loci by allelic replacement, as described in (Diebold-Durand et al., 2019). Strains were selected on SMG-agar plates under appropriate antibiotic selection. Genotypes were verified for single colony isolates by PCR and Sanger sequencing as required. A list of strains and genotypes are given in Supplementary Table 1. An assignment of strains to figure panel is listed in Supplementary Table 2.

### Viability assessment by dilution spotting

Cultures were inoculated in SMG medium and grown for 8 hrs at 37 °C under constant agitation. Cultures were diluted 1:9 in series. Dilutions of 9^2^ and 9^5^ were spotted on SMG-agar (SMM glucose glutamate tryptophane) and ONA (Oxoid nutrient agar) plates and grown at 37 °C. Colony growth was documented by imaging after 16 hrs for ONA plates and 24 hrs for SMG plates.

### Chromosome conformation capture coupled with deep sequencing (3C-seq)

3C-seq was performed essentially as described in (Anchimiuk *et al*., 2021).

#### Sample collection

Cultures were grown in SMG at 37 °C in mid-exponential phase (OD600 = 0.02-0.03) and fixed with formaldehyde (3% final concentration) for 30 min at RT and 30 min at 4 °C. The formaldehyde crosslinking was quenched by 30 min incubation with 0.25 M glycine at 4 °C. Finally, the cells were pelleted by filtration, washed with fresh SMG and frozen in liquid nitrogen for storage at −80 °C.

#### Cell pellet processing

To lyse the cells, the 3C cell pellets were resuspended in 600 μl 1× TE (Sigma) supplemented with 4 μl of Ready-lyze lysozyme (35 U/μl, Tebu Bio). After 20 min incubation at RT, SDS was added to a final concentration of 0.5% and incubated for additional 10 min. 50 μl of lysed cells were aliquoted to 8 tubes containing 450 μl of digestion mix (1× NEB 1 buffer, 1% triton X-100, and 100 U HpaII enzyme [NEB]) and incubated at 37 °C for 3 hours with constant shaking. Fragmented DNA was collected by centrifugation, resuspended in 800 μl 1× TE and diluted into 4 tubes containing 8 ml of ligation mix (1× ligation buffer: 50 mM Tris-HCl, 10 mM MgCl2, 10 mM DTT, 1 mM ATP, 0.1 mg/ml BSA, 125 U T4 DNA ligase 5 U/ml) and incubated at 16°C for 4 hours. Proximity ligation reaction was followed by O/N decrosslinking at 65°C in the presence of 250 μg/ml proteinase K (Eurobio) and 5 mM EDTA (Sigma).

#### DNA purification

To purify the DNA, isopropanol precipitation was performed. Each sample was mixed with 1 volume of isopropanol and 0.1 volume of 3M NaOAc (pH 5.2, Sigma) and incubated at −80 °C for 1 hour. The DNA was collected by centrifugation and resuspended in 400 μl 1× TE at 30 °C for 20 min. Next, phenol-chloroform-isoamyl alcohol extraction was performed, followed by final DNA precipitation using 1.5 volume of cold 100% EtOH in the presence of 0.1 volume of 3M NaOAc at −80 °C for 30 min. Collected pellets were resuspended in 30 μl 1 x TE and incubated with RNaseA at 37 °C for 30 min. All tubes belonging to the same sample were pooled and the resulting 3C library was quantified on gel using ImageJ.

#### Library preparation and sequencing

All 3C libraries were adjusted to 1 μg for library preparation. Each 3C library volume was adjusted to 130 μl and sonicated using Covaris S220 following 500 bp target size recommendations from the manufacturer. Fragmented DNA was purified using Qiagen PCR purification kit, eluted in 40 μl EB and quantified using the NanoDrop. Custom-made adapters were used to prepare the libraries for paired-end Illumina sequencing using ^~^1 μg of DNA as an input. Adapter ligation was performed for 4 hours at RT, followed by an inactivation step at 65 °C for 20 min. DNA was purified with 0.75× AMPure beads and 3 μl were used for 50 μl PCR reaction (12 cycles). Amplified libraries were purified on Qiagen columns and pair-end sequenced on an Illumina platform (HiSeq4000 or NextSeq).

#### Processing of PE reads and generation of contact maps

A custom script was used to demultiplex the sequencing data. Prinseq was used to clean the data prior to processing it following the steps described at https://github.com/axelcournac/3C_tutorial.

Briefly, each mate was mapped to the reference genome (NC_000964.3) using bowtie2 in very-sensitive-local mode. Next, data was sorted and both mates merged. The reads of mapping quality above 30 were filtered out and assigned to a restriction fragment. Uninformative events including recircularization on itself (loops), uncut fragments, and re-ligations in original orientation were discarded. Only pairs of reads corresponding to long-range interactions were used for generation of contact maps (between 5 and 8% of all reads). A bin size of 10 kb was used. Contact maps were normalized through the sequential component normalization procedure (SCN). Subsequent visualization was done using MATLAB (R2019b). To facilitate visualization of the contact maps, first the log10 and then a Gaussian filter (H = 1) were applied to smooth the image.

### Live cell imaging

Cells were grown in SMG to OD600 = 0.04. 2 ml culture volumes were spun down at 8000 rcf for 2 minute at RT. Supernatant was removed, cells were resuspended in 10 μl PBS. 0.5 μl cell suspension were spotted onto homemade agarose microscopy slides. Images were acquired using a Leica DMi8 microscope with a sCMOS DFC9000 (Leica) camera, a SOLA light engine (Lumencor) and a ×100/1.40 oil-immersion objective. Images were acquired with 600 ms exposure at 470 nm excitation, 520 nm emission. Images were processed using LasX v.3.3.0.16799 (Leica).

### Chromatin-Immunoprecipitation coupled to quantitative PCR (ChIP-qPCR)

ChIP samples were prepared as described previously (Bürmann *et al*., 2017). Cultures were grown in 200 ml volumes SMG at 37°C. Cells were grown to mid-exponential phase (OD600=0.02-0.03) and fixed by incubation for 30 minutes with 1/10 [v/v] of buffer F (50 mM Tris-HCl pH 7.4, 100 mM NaCl, 0.5 mM EGTA pH 8.0, 1 mM EDTA pH 8.0, 10% [w/v] formaldehyde). Cells were harvested by filtration and washed in cold PBS. OD600 values of samples were normalized to 2 and resuspended in TSEMS (50 mM Tris pH 7.4, 50 mM NaCl, 10 mM EDTA pH 8.0, 0.5 M sucrose and PIC (Sigma)) supplemented with 6 mg/ml chicken egg white lysozyme (Sigma). Samples were incubated at 37°C for 30 minutes under shaking. Resulting protoplasts were harvested by centrifugation, washed in 2 ml TSEMS, resuspended in ml TSEMS and split into 3 aliquots of equivalent volume before pelleting and flash freezing.

Samples were resuspended in 2 ml of buffer L (50 mM HEPES-KOH pH 7.5, 140 mM NaCl, 1 mM EDTA pH 8.0, 1% [v/v] Triton X-100, 0.1% [w/v] Na-deoxycholate, 0.1 mg/ml RNaseA and PIC (Sigma)), transferred to 5 ml round-bottom tubes and sonicated three times for 20 seconds using a Bandelin Sonoplus with a MS72 tip (90% pulse and 35% power output). Suspensions were centrifuged for 10 minutes at 21 krcf at 4°C. Samples were split into 200 μl Input material and 800 μl IP material.

Antibody serum was incubated with equivalent volumes of Dynabeads Protein G suspension (Invitrogen) for 2 hours at 4°C under gentle agitation. Beads were washed in 1 ml Buffer L directly prior to use and resuspended as 50 μL aliquots of 30 mg/mL. IP material was mixed with these 50 μl aliquots and incubated at 4°C for 2 hours under rotation. Bound material was subsequently washed by 1 ml washes with buffer L, L5 buffer L containing 500 mM NaCl), buffer W (10 mM Tris-HCl pH 8.0, 250 mM LiCl, 0.5% [v/v] NP-40, 0.5% [w/v] Na-Deoxycholate, 1 mM EDTA pH 8.0) and buffer TE (10 mM Tris-HCl pH 8.0, 1 mM EDTA pH 8.0). Beads were resuspended in 520 μL TES (50 mM Tris-HCl pH 8.0, 10 mM EDTA pH 8.0, 1% (w/v) SDS). Input material was supplemented with 300 μl TES and 20 μl 10% SDS. Tubes were incubated at 65°C over-night under vigorous shaking.

DNA was purified using Phenol-chloroform extraction by adding and thoroughly mixing first with 500 μl phenol equilibrated with buffer (10 mM Tris-HCl pH 8.0, 1 mM EDTA). Phases were separated by centrifugation for 10 min at 21 krcf. 450 μL supernatant were subsequently mixed with and again separated from 450 μL chloroform. 400 μL supernatant were taken off, mixed with 1.2 μl GlycoBlue (Invitrogen), 30 μl of 3 M Na-Acetate (pH 5.2) and 1 ml ethanol (filtered). Samples were incubated at −20°C for at least 30 minutes. DNA was precipitated by centrifugation at room temperature with 21 krcf for 10 minutes. DNA was resuspended in 100 μL volumes PB (Qiagen) and dissolved by incubation at 55°C under vigorous shaking for 10 minutes. Samples were subsequently purified using a PCR purification kit (Qiagen) as per protocol and eluted in 50 μl elution buffer.

Samples were diluted to 1:10 for IP and 1:100 for Input material. Reaction for qPCR were prepared by mixing 4 μl diluted samples with 5 μl 2x 5 μl Takyon SYBR MasterMix and 1 μl qPCR primer mixture (3 μ M). A list of qPCR primers in given in Supplementary Table 3. Samples were run in a Rotor-Gene Q (Qiagen) and analyzed using PCR miner (Zhao and Fernald, 2005).

### *In vivo* Cys-Cys and Lys-Cys cross-linking

Cross-linking was performed das described in (Soh *et al*., 2019). Cultures were grown in SMG to OD600 of 0.03-0.04 at 37°C. Cells were mixed with ice and harvested by centrifugation. Samples handling and preparation was done on ice and cold at every step. Cells were washed in PBSG (PBS with 0.1% [v/v] glycerol). Samples were resuspended in 1 ml PBSG. 1.25 OD600 equivalents were taken and pelleted by centrifugation. Pellets were resuspended in 30 μl PBSG. Cross-linking agent (SMCC (Thermo) or BMOE (Thermo) were added to 0.5 mM final concentration and mixed by vortexing. Reactions were incubated on ice for 10 minutes and then quenched by addition of 0.5 mM final concentration 2-Mercaptoethanol with subsequent incubation for 2 minutes. Samples were supplemented with additives at the indicated final concentrations: Benzonase (750 U/ml; Sigma), 5 μM HaloTag-TMR ligand (Promega), Ready-Lyse Lysozyme (47 U/μl; Epicentre), and 1× PIC (Sigma-Aldrich). Samples were incubated at 37°C for 30 minutes under light protection. Samples were supplemented with LPS loading dye and denatured at 70°C for 5 minutes. Samples were run on 3-8% Tris-Acetate gels (Invitrogen) at constant power output of 35 mA at 4°C. In-gel fluorescence was imaged using an Amersham Typhoon (GE Healthcare) with a Cy3 DIGE filter. Quantification was done using ImageQuant (GE Healthcare).

### Structure prediction by AlphaFold-Multimer

Predictions were performed using the Colab notebook (dpmd.ai/alphafold-colab) (Evans *et al*., 2021; Jumper et al., 2021). The input sequences are denoted in the figure legends.

